# Semi-supervised detection of natural selection with positive-unlabeled learning

**DOI:** 10.1101/2025.08.15.670602

**Authors:** Sandipan Paul Arnab, Andre Luiz Campelo dos Santos, Matteo Fumagalli, Michael DeGiorgio

## Abstract

Identifying genomic regions shaped by natural selection is a central goal in evolutionary genomics. Existing machine learning methods for this task are typically trained using simulated genomic data labeled according to specific evolutionary scenarios. While effective in controlled settings, these models are limited by their reliance on explicit class labels. They can only detect the specific processes they were trained to recognize, making it difficult to interpret predictions for regions influenced by other evolutionary forces. This limitation is especially problematic when analyzing empirical genomes shaped by a mixture of adaptive, demographic, and ecological factors. One-vs.-rest strategies offer a potential alternative, but suffer from the inherent complexity of modeling all other evolutionary and demographic processes as a catch-all “rest” class. Here, we explore positive-unlabeled learning as a more flexible framework for detection of adaptive events. Positive-unlabeled learning is a semi-supervised approach that permits identification of samples of a target class using only positive labels and an unlabeled background, without requiring explicit modeling of negative samples. To assess the utility of this approach, we focus on a binary classification setting for detecting selective sweeps (positive samples) arising from positive natural selection against a mixed background composed of both unlabeled sweeps and neutrally-evolving regions. To accomplish this goal, we introduce *PULSe*, a method that employs only a set of labeled sweep observations for training while treating all remaining data as unlabeled. By avoiding assumptions about the composition of the background, *PULSe* enables robust sweep discovery in realistic genomic landscapes. We systematically evaluate its performance across a range of demographic, adaptive, and confounding contexts, including domain shift arising from misspecified demographic models, and find that *PULSe* delivers high performance and generalizability. To demonstrate a practical application of *PULSe*, we analyzed European and Bengali in Bangladesh human genomes, treating the empirical genomes as unlabeled sets, and recapitulating several previously-identified sweep candidates. Our results show that *PULSe* provides a powerful and versatile alternative for detecting adaptive regions, with the potential to generalize across a range of genomic landscapes.

## Introduction

Detecting genomic regions influenced by adaptive evolution is critical for uncovering how organisms respond to their environments and for identifying loci that contribute to fitness-related traits [Pigllucci, 1996, Ellegren and Sheldon, 2008]. Genomic regions influenced by adaptive processes often bear distinctive signals, including altered variation, or skewed allele frequency distributions that reflect their departure from neutrality [Lewontin and Krakauer, 1973, Günther and Coop, 2013]. Detecting these regions offers a powerful means to infer the genetic basis of adaptation, illuminate species histories, and identify functionally important loci [Pardo-Diaz et al., 2015]. However, the multiplicity of adaptive modes makes comprehensive identification difficult. Each mode can leave different, sometimes overlapping footprints in genetic data [Oleksyk et al., 2010], often entangled with confounding demographic effects [Saccheri and Hanski, 2006, Byars et al., 2010]. While empirical-based approaches [Voight et al., 2006] have the capacity to correct for demographic effects, they do not provide a framework for inferring the proportions of sites under adaptive evolution. The additional challenge lies not only in distinguishing adaptive from neutral regions, but in building models that can generalize beyond any single evolutionary process.

Modern machine learning methods have shown promise in this domain by learning discriminative patterns across many features simultaneously, whether derived from summary statistics or raw genomic alignments [Sheehan and Song, 2016, Schrider and Kern, 2017, Sugden et al., 2018, Flagel et al., 2019, Torada et al., 2019, Mughal et al., 2020, Gower et al., 2021, Lauterbur et al., 2023, Whitehouse and Schrider, 2023, Ahlquist et al., 2023, Cecil and Sugden, 2023, Arnab et al., 2025]. These tools can flexibly capture relationships between input data and evolutionary outcomes, providing an appealing alternative to single-test statistics. Some recent approaches have adopted deep learning architectures, such as convolutional neural networks, to classify genomic windows by selection type or evolutionary history, often reporting superior performance over traditional methods [Kern and Schrider, 2018, Hejase et al., 2022, Arnab et al., 2023]. Despite these advances, most machine learning approaches in population genomics remain constrained by their dependence on simulated, explicitly labeled training data. Typically, models are trained to distinguish one or more types of selection from neutral evolution, relying on labels derived from simulations under predefined models. As a result, they tend to learn only the specific patterns they are trained to recognize. This limitation constrains their utility when applied to empirical data, where the true evolutionary history, which includes both demographic history and the full spectrum of adaptive processes, may be far more complex or entirely unmodeled.

The core limitation lies in how the “negative class” is defined. In many supervised machine learning pipelines, the model is trained to classify between a selective process of interest and a negative (background) class, generally comprising neutrality. Yet real genomes are shaped by a mixture of nonadaptive forces like mutation, migration, drift, population size changes, and demographic shifts, as well as adaptive forces acting through various mechanisms [Kimura, 1979, Slatkin, 1987, Barton and Charlesworth, 1998, Lynch, 2010]. Attempting to simulate all possible evolutionary processes affecting the background class would involve substantial efforts in simulation and modeling while not being comprehensive enough, as we never know the full array of forces acting on empirical genomes. Even with expansive simulation sets, a classifier trained on specific assumptions may fail when applied to data generated under different histories. This mismatch, which can be characterized as domain shift [Ganin et al., 2016, Luo et al., 2019, Stacke et al., 2020, Zhang et al., 2021], can degrade model accuracy and lead to overconfident and incorrect predictions. One-vs-rest classifiers [Xu, 2011, Bali and Mansotra, 2021], which can treat all other forms of adaptive and non-adaptive processes except for the targeted mode of adaptation as a generic “rest” class, inherit this limitation and may confuse unknown selection modes with noise. Consequently, there is a need for detection frameworks that do not rely on explicitly modeling all possible evolutionary alternatives.

To address this issue, we introduce a strategy based on positive-unlabeled (PU) learning, a form of semi-supervised learning designed for classification problems where only a subset of positive samples are labeled, and no explicitly labeled negatives are available [Elkan and Noto, 2008]. PU learning has been successfully applied in domains such as fraud detection and medical diagnosis [Chen et al., 2020, Vinay et al., 2022], where true negatives are hard to define or curate. Its core assumption is that labeled positives represent one class of interest, while the unlabeled data contains a mixture of positives and negatives. The goal is to distinguish positives from this unlabeled background without ever specifying what a negative should look like. In many real-world PU learning applications, the unlabeled set is highly imbalanced, with positives representing only a small fraction of the total [Su et al., 2021]. This assumption maps naturally to evolutionary genomics, where we may have only a few high-confidence examples of regions under selection, while the vast majority of the genome is expected to be neutrally evolving or shaped by other nonadaptive processes. PU learning thus offers a principled way to bypass the need for explicit negative labels, allowing models to learn what distinguishes known adaptive regions from the rest of the genome, regardless of the composition of the background.

Here, we present the method *PULSe* (*P* ositive-*U* nlabeled *L*earning for *Se*lection detection), which applies this framework to the task of identifying genomic regions shaped by adaptation. *PULSe* requires only a labeled set of adaptive samples and treats the remaining data as unlabeled, without making assumptions about the evolutionary processes shaping that background. While the method is intended to generalize to a wide range of adaptive scenarios, we use selective sweeps as a benchmark for validation. A selective sweep is a phenomenon that is introduced by positive natural selection. When a beneficial genetic mutation quickly spreads through a population, it reduces genetic variation in nearby regions of the genome [Przeworski, 2002, Hermisson and Pennings, 2005, Stephan, 2016, 2019]. Sweeps are a convenient test case because they produce well-characterized genomic signals of adaptation and can be readily simulated using widely available population genetic simulation software [Ewing and Hermisson, 2010, Kern and Schrider, 2016, Haller and Messer, 2019]. We assess the ability of *PULSe* to detect sweep regions when trained only on positive samples, in the presence of challenging confounders such as demographic model misspecification, and background selection.

We evaluate *PULSe* across an extensive array of simulation experiments, comparing its performance to that of fully supervised models. These evaluations include both in-domain and out-of-domain testing to assess the robustness of the method to domain shift. By design, *PULSe* is modular and compatible with a variety of feature generation protocols and classifier algorithms, making it adaptable to different organismal systems and data types. *PULSe* is also user-friendly and can be seamlessly integrated with common simulators, provided that the simulations can be exported in *ms* [Hudson, 2002] or Variant Call Format [Danecek et al., 2011] files. As a demonstration of its practical utility, we apply *PULSe* to human polymorphism data from Europeans and Bengali in Bangladesh, treating the genomes as unlabeled backgrounds and using simulated sweeps as positive training samples. The model recovers several well-characterized sweep candidates previously identified in empirical scans and flags additional regions of interest for further investigation. These results suggest that *PULSe* can serve as a flexible tool for detecting regions shaped by selection in empirical genomes.

## Results

### Modeling description

We utilized the coalescent simulator discoal [Kern and Schrider, 2016] to generate neutral and sweep replicates under a nonequilibrium demographic history estimated for European (CEU) humans [Tennessen et al., 2012]. This demographic model includes a recent severe population bottleneck, capturing the complex evolutionary dynamics of this population. To simulate sweeps, we considered per-generation selection coefficients (*s*) ranging from 0.005 to 0.1, initial beneficial allele frequencies (*f*) spanning from 1*/*(2*N_e_*) to 0.2, and fixation times (*τ*) varying from 0 to 2,000 generations before sampling. Additionally, both neutral and sweep replicates were generated with per-site per-generation mutation rates (*µ*) ranging from 2.21 × 10^−9^ to 2.21 × 10^−8^, and per-site per-generation recombination rates (*r*) from 0 to 3 × 10^−8^, with a mean rate of 10^−8^. These parameters were chosen to comprehensively capture the evolutionary forces acting on the CEU population while ensuring that a wide spectrum of selective and neutral genetic patterns was examined. For specific details about the simulations, refer to the *Simulation protocol* subsection of the *Materials and Methods*.

To further investigate the impact of demographic assumptions, we also generated neutral and sweep replicates based on a constant-size demographic model while maintaining the same genetic and adaptive parameter settings as in the CEU simulations. The only exception was the diploid effective population size, which we varied across a range from 3,000 to 30,000 in increments of 250. Given the breadth of selection strengths, beneficial allele frequencies, and fixation times explored, along with the inherent variation in mutation and recombination rates, distinguishing between neutral and selective scenarios remains challenging. This difficulty is further compounded by the potential for substantial overlap in distributions of genetic variation between sweeps and neutrality, emphasizing the necessity of robust and adaptable detection methods.

To construct the input for *PULSe*, we first generated images [Arnab et al., 2025] from the CEU and constant-size simulated replicates. These images were created by transforming minor allele count data into sorted representations across short, overlapping windows. The image generation process (Figure 1A) from simulated data is detailed in the *Image generation* subsection of the *Materials and Methods*. To extract informative features from these images, we applied histogram of oriented gradients (HOG) [Freeman and Roth, 1995], which captures local gradient orientations and edge structures (Figure 1B), effectively encoding spatial patterns indicative of changes in genomic diversity (see *Histogram of oriented gradients* subsection of the *Materials and Methods* for details). We then used these extracted HOG features in a positive-unlabeled (PU) learning framework (Figure 1C), where logistic regression served as the base classifier (see *Positive-unlabeled learning algorithm* subsection of the *Materials and Methods* for details).

**Figure 1:**
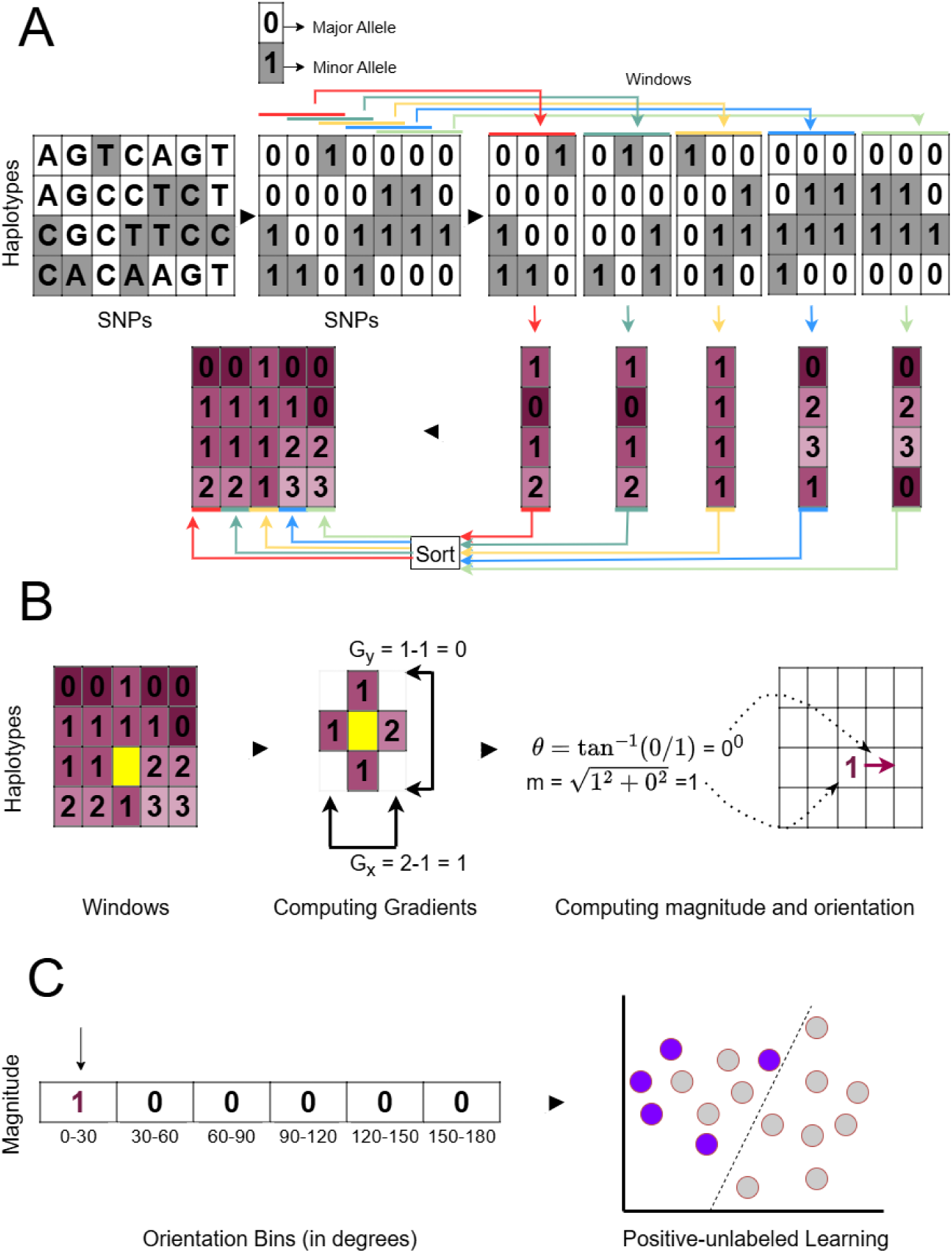
Depiction of the *PULSe* model. (A) As described in the *Image generation* subsection of the *Materials and Methods*, for a sample of haplotypes (rows) across a set of SNPs (columns), the major and minor alleles at each SNP are coded as zero and one, respectively. From this processed alignment, the number of minor alleles for each haplotype is counted within a window of fixed size (here of three SNPs) and window locations are shifted by a specific stride (here of one SNP). The minor allele counts for each window are then sorted, such that the top row has the smallest value and the bottom row the largest value for a given window. A matrix is computed based on a specified number of consecutive windows (here five windows), which we consider as an input image. (B) For a given pixel in this image, gradient magnitude *m* and orientation *θ* are calculated based on the differences in pixel intensities along the horizontal (*G_x_*) and vertical (*G_y_*) directions. (C) These gradient magnitudes across the image are then accumulated into orientation bins, illustrated here with one such gradient, to build a histogram of oriented gradients vector that is used as input to a positive-unlabeled learning framework.

Unlike traditional supervised learning, PU learning does not require a fully labeled training dataset. Instead, it relies on a set of positively labeled samples (sweep regions in this case) while treating the remaining data as an unlabeled mixture of negative (non-sweep regions) and potentially undetected positive (sweep regions) samples. For simulated testing, we designed two scenarios to evaluate *PULSe*. In the first scenario, we used 10,000 CEU sweep replicates as the labeled set. We then used 20,000 CEU simulations as the unlabeled set, introducing varying degrees of imbalance between sweep and neutral observations within this set. Because most of the genome is expected to evolve neutrally [Kimura et al., 1968, Jensen et al., 2019], the majority of the unlabeled set consisted of CEU neutral replicates, with the specific proportions of sweep and neutral replicates detailed in the *Model performance as function of unlabeled set class imbalance* subsection below. The second scenario followed the same structure but used 10,000 constant population size sweeps as the labeled set while keeping the same CEU unlabeled set as in the first scenario. We respectively denote these two scenarios as *CEU-CEU* (domain matched) and *Constant-CEU* (domain mismatched) for ease of reference. These experimental setups allow us to assess the robustness of PU learning in detecting selection in the CEU demographic context.

Heatmaps depicting mean images for the sweep and neutral classes in both the CEU and constant-size datasets reveal distinct patterns associated with positive selection. In contrast to the neutral image, the sweep image exhibits a dark vertical segment in the central columns, representing a loss of genetic variation due to high-frequency haplotypes (Figure 2). This pattern arises because major alleles occur at high frequency near the center of the sweep replicates, reflecting the impact of selection on linked variation. By presenting mean images for both CEU and constant population size sweep and neutral replicates, we illustrate the consistency of these patterns across different demographic contexts.

**Figure 2:**
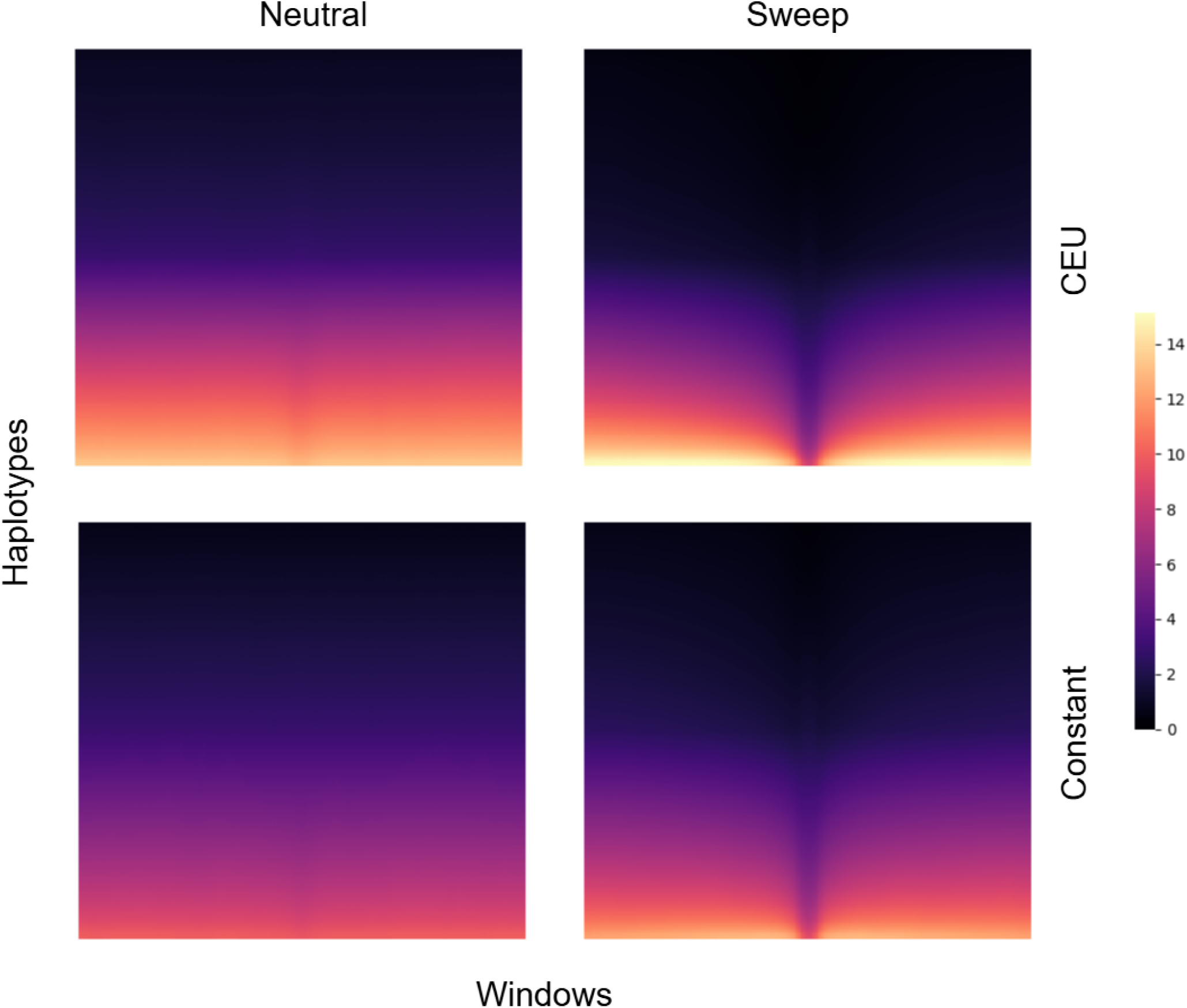
Heatmaps depicting input images of size 198 × 238, averaged across 10,000 neutral and 10,000 sweeps replicates simulated under either the CEU (top row) or constant size demographic history (bottom row). Input images are processed as in the *Image generation* subsection of the *Materials and Methods*. Rows of the images represent locally sorted minor allele counts within haplotype windows, whereas columns represent genomic windows of 25 contiguous SNPs within a haplotype, with an equal number of windows flanking the center of a simulated genomic region. The colorbar indicates the number of minor alleles within the haplotype window, with darker colors representing higher numbers of major alleles and brighter colors higher numbers of minor alleles.

In addition, we thoroughly assessed various HOG parameters and pre- and post-processing pipelines, with full details provided in the *Choosing the best feature extraction pipeline* subsection of the *Materials and Methods*. After extensive testing, we identified the *P1* and *P2* pipelines as performing best in the domain matched and mismatched scenarios, respectively, leading to the development of the *PULSe*[*P1*] and *PULSe*[*P2*] models (see *Choosing the best feature extraction pipeline subsection* for details of these pipelines). To evaluate the effectiveness of different feature extraction pipelines, we used the area under the precision-recall curve (AUPRC) as our performance metric. This choice was motivated by the particular suitability of the AUPRC for imbalanced datasets, where it better reflects the ability of a model to identify the positive (minority) class by emphasizing precision (positive predictive value) and recall (true positive rate) over overall accuracy [Saito and Rehmsmeier, 2015]. For context, in datasets with only 10% positive samples, a baseline AUPRC expected from random guessing would fall around 0.10 [Saito and Rehmsmeier, 2015]. To complement AUPRC, we also evaluated the Matthews correlation coefficient (MCC), which is particularly robust for binary classification in imbalanced and partially labeled datasets [Boughorbel et al., 2017, Chicco and Jurman, 2020]. MCC offers a single balanced metric that incorporates both true and false predictions across classes (class-specific detection rates), complementing AUPRC, which primarily reflects performance on the minority positive class.

### Model performance as function of unlabeled set class imbalance

To examine how the degree of class imbalance of the unlabeled set affects model performance, we conducted experiments while fixing the number of labeled and unlabeled samples to 10,000 each. We varied the proportion of positive samples in the unlabeled set, testing levels of 2%, 4%, 6%, 8%, and 10%. Despite these changes, the neutral detection rate remained virtually unchanged across all tested conditions for both the *CEU-CEU* (ranging from 95.5% to 95.9%) and *Constant-CEU* (ranging from 88.3% to 88.4%) scenarios (Figure S1). Furthermore, no distinct performance gap was observed between *PULSe*[*P1*] and *PULSe*[*P2*] in either scenario, suggesting that the proportion of positive samples in the unlabeled set does not significantly impact model behavior under these conditions.

Sweep detection rate, however, was heavily affected by the degree of class imbalance of the unlabeled set. As the proportion of positive samples increased, sweep detection rate dropped significantly, suggesting that at their present configurations, *PULSe* models tend to detect adaptive regions better with higher degrees of class imbalance. In the *CEU-CEU* scenario, the detection rate for *PULSe*[*P1*] decreased by 6.6%, whereas *PULSe*[*P2*] experienced a slightly larger drop of 7.3%. The effect was even more pronounced in the *Constant-CEU* scenario, where both models saw a decline of approximately 10.4%.

However, as the proportion of positive samples increased, the AUPRC noticeably improved. In the *CEU-CEU* scenario, AUPRC increased by 0.215 for *PULSe*[*P1*] and 0.220 for *PULSe*[*P2*], whereas for the *Constant-CEU* scenario, the increase was 0.107 for *PULSe*[*P1*] and 0.136 for *PULSe*[*P2*]. This discrepancy between the decline in sweep detection rate and the rise in AUPRC suggests that the models may be miscalibrated with respect to class proportions [McDermott et al., 2024]. It is possible that the model may be shifting decision thresholds in response to the higher proportion of positive samples, leading to higher AUPRC while simultaneously missing sweep signals. Additionally, though MCC shows a similar upward trend as the proportion of positive samples increases, its growth is not as sharp. In the *CEU-CEU* scenario, the MCC of *PULSe*[*P1*] increases by 0.103 and by 0.127 for *PULSe*[*P2*], whereas in the *Constant-CEU* scenario, the increases are 0.017 and 0.013, respectively.

### Effect of the ratio of labeled to unlabeled samples

To evaluate how varying the ratio of labeled to unlabeled samples impacts *PULSe* model performance, we examined six test cases in which the number of labeled samples was fixed at 10,000, and the number of unlabeled samples was set to 10,000, 12,000, 14,000, 16,000, 18,000, and 20,000 (Figure S2). In each scenario, 10% of the unlabeled samples consist of sweeps, whereas the remainder are neutral replicates.

*PULSe*[*P1*] under the *CEU-CEU* setting shows AUPRC values increasing from 0.7197 to 0.7669 as the number of unlabeled samples increases, whereas in the *Constant-CEU* scenario it exhibits a smaller improvement in AUPRC, increasing from 0.3321 to 0.3411. On the other hand, though *PULSe*[*P2*] under the *CEU-CEU* setting showcases an increase in AUPRC values from 0.7106 to 0.7592, we notice a drop from 0.3794 to 0.3547 in the *Constant-CEU* scenario. Though the *P1* pipeline exhibits an upward trend in AUPRC and *P2* a downward trend with increasing numbers of unlabeled samples in the *Constant-CEU* scenario, *P2* consistently achieves higher AUPRC than *P1* across all test cases.

We observe a similar trend in the MCC values. For instance, *PULSe*[*P1*] under the *CEU-CEU* scenario shows MCC values increasing from 0.6286 to 0.6655 as a function of the number of unlabeled samples, whereas in the *Constant-CEU* setting it exhibits a rise from 0.2941 to 0.3112. However, though *PULSe*[*P2*] under the *CEU-CEU* scenario demonstrates an increase in MCC values from 0.6253 to 0.6733, in the *Constant-CEU* setting, MCC drops from 0.3368 to 0.3205. Just like AUPRC, MCC follows the same pattern, with *P2* outperforming *P1* in every *Constant-CEU* test case despite the opposite directional trends.

Across both scenarios, both *PULSe* model variants consistently exhibit increasing neutral detection accuracy as the number of unlabeled samples increases. In the *CEU-CEU* scenario, *PULSe*[*P1*] neutral detection accuracy rises from 94.42% with 10,000 unlabeled samples to 97.57% with 20,000 unlabeled samples, whereas *PULSe*[*P2*] demonstrates a similar improvement, increasing from 93.96 to 97.61%. The *Constant-CEU* setting presents a similar trend of increase in neutral detection accuracies, with *PULSe*[*P1*] rising from 86.86 to 93.88% and *PULSe*[*P2*] from 87.14 to 93.74% as a function of the number of unlabeled samples.

Sweep detection accuracy on the other hand, decreases consistently as the number of unlabeled samples grows. In the *CEU-CEU* scenario, *PULSe*[*P1*] sweep detection accuracy declines from 74.90 to 64.85%, with *PULSe*[*P2*] harboring a slightly larger decrease from 76.30 to 64.70%, indicating that the ability of the model to detect the minority sweep class diminishes with increased unlabeled to labeled ratio. The *Constant-CEU* scenario results further attest to this trend, with *PULSe*[*P1*] experiencing a drop in sweep accuracy from 49.80 to 35.95% and *PULSe*[*P2*] falling from 55.10 to 37.40%.

### Improving detection performance with model calibration

To assess the reliability of the *PULSe* class probability estimates, we examined the calibration of our classifiers. Because calibration curves are sensitive to class imbalance [Zadrozny and Elkan, 2002, Niculescu-Mizil and Caruana, 2005], we constructed our calibration test set by selecting 50% of the sweep samples uniformly at random from the labeled set along with an equal number of random samples from the unlabeled set. We find that for both *PULSe*[*P1*] and *PULSe*[*P2*] models in both the *CEU-CEU* and *Constant-CEU* scenarios, the calibration curves deviate markedly from the ideal *y* = *x* line, indicating significant miscalibration of the predicted probabilities (Figure S3). Performing calibration within the PU learning framework is not straightforward because the labeled set contains only positive samples, and in empirical applications we cannot assume the true class labels for the unlabeled set. To address this challenge, we devised a novel approach for calibration of *PULSe* models. Specifically, we chose to apply isotonic regression [Zadrozny and Elkan, 2002] for calibration, though we acknowledge that Platt scaling can also be used, which as a parametric method, requires learning two parameters through training and validation, and can help reduce overfitting [Platt et al., 1999]. In our approach, we use the probability estimates of known sweeps from the labeled set as the positive calibration targets. For the negative class, we select original samples from the unlabeled set whose raw sweep probabilities fall below various thresholds. This strategy allows us to construct a calibration set that better approximates the balanced distribution required for effective calibration.

To thoroughly evaluate the effectiveness of our calibration approach, we conducted experiments across multiple configurations. We fixed the number of positive samples in the labeled set to 10,000, while varying the number of unlabeled samples between 10,000 and 20,000, and choosing the proportion of sweeps in the unlabeled set to either 5% or 10%. Additionally, we explored an array of positive class probability thresholds for selecting negative samples from the unlabeled set, with these cutoff values drawn from 0.2, 0.3, 0.4, 0.5, 0.6, or 0.7. Though a positive class probability threshold of 0.5 would seem the most natural choice, we examined this broader range because, particularly in the *Constant-CEU* scenario, both *PULSe* models were notably weak in detecting sweeps.

Our results show that regardless of the configuration, the calibration approach improved the calibration of the models (Figure S3). However, several general patterns emerged. Consequently, we could not identify a single pipeline that performs optimally across both scenarios. Rather, the findings suggest that in cases of domain match, the simpler *P1* pipeline performs better, whereas in cases of domain mismatch, the more complex *P2* pipeline is more effective. This difference may arise because, in a domain matched setting, the additional processing in *P2* introduces unnecessary complexity that can dilute discriminative features already well-aligned between training and test data. We hypothesize that with *P2*, histogram equalization (see *Choosing the best feature extraction pipeline* subsection of the *Materials and Methods* for details) helps preemptively mitigate the effect of domain shift, while the higher number of orientation bins is needed to capture more subtle changes in gradients under a domain mismatched setup. Additionally, when examining *PULSe*[*P1*] in the *CEU-CEU* scenario and *PULSe*[*P2*] in the *Constant-CEU* setting, we observed that the calibration performance improves as the number of unlabeled samples increases, regardless of the proportion of sweeps in the unlabeled set (Figure S3). This behavior is expected, as isotonic regression is less prone to overfitting with a larger number of training samples [Niculescu-Mizil and Caruana, 2005].

A positive class probability threshold of 0.3 generally resulted in the highest post-calibration AUPRC across both *PULSe* models and demographic scenarios (Figures S4 and S5). Accordingly, we selected 0.3 as the calibration threshold for our empirical analyses. Comparing post-calibration MCC against pre-calibration MCC on the unlabeled set containing 10,000 samples and percentage of sweep samples set at 10% shows a small increase of 0.066 for *PULSe*[*P1*] on *CEU-CEU* (Figure S6 vs. Figure S4), and a substantial increase of 0.1898 for *PULSe*[*P2*] on *Constant-CEU* (Figure S6 vs. Figure S5).

### Hyperparameter tuning for the base classifier

So far we have used an unpenalized logistic regression model as the base classifier in *PULSe*. Though this choice provided a simple starting point to judge the viability of the PU learning process with logistic regression as the base classifier, it can be prone to overfitting as HOG features derived from *PULSe* images can exhibit high correlation due to the spatial symmetry of the images. To improve generalization and enhance robustness across settings, we introduced elastic net regularization when fitting the logistic regression classifier, which combines both *L*_1_- and *L*_2_-norm penalties [Zou and Hastie, 2005, Hastie et al., 2009]. Elastic net regularization encourages the model to use only a few important features while keeping the solution stable and less sensitive to noisy or correlated features.

We performed a grid search to identify the best combination of regularization parameters based on the highest AUPRC on the unlabeled set. Specifically, we varied the mixing parameter *α* ∈ {0.0, 0.1, …, 1.0}, which controls the balance between *L*_1_- and *L*_2_-norm penalties with *α* = 1 corresponding to pure *L*_1_-norm regularization and *α* = 0 to pure *L*_2_-norm regularization, and the regularization strength *λ* ∈ {10^−6^, 10^−5^, …, 10^5^}. For each combination, we trained the classifier on a dataset consisting of 10,000 labeled sweep replicates and 10,000 unlabeled samples, with the percentage of sweeps in the unlabeled dataset fixed at 10%.

For *PULSe*[*P1*] evaluated under the matched *CEU–CEU* demographic scenario, the optimal classifier achieved an AUPRC of 0.727, and an MCC of 0.6576 using *α* = 0.6 and *λ* = 1. In contrast, under the domain mismatched *Constant–CEU* setting, *PULSe*[*P2*] performed best with *α* = 0.2 and *λ* = 0.01 (Figure 3). Though overall performance is still lower under domain shift, the use of regularization along with the calibration mechanism (see *Improving detection performance with model calibration* subsection above) helped reduce the performance gap between matched and mismatched settings. While sweep detection accuracy was comparable across models, the key difference lies in neutral detection as *PULSe*[*P1*], trained and tested under a matched demographic scenario, achieved a 12.4% higher neutral detection accuracy than *PULSe*[*P2*], which operated under domain mismatch. *PULSe*[*P2*] thus reports substantially lower AUPRC (0.6531) and MCC (0.4907) relative to *PULSe*[*P1*] on domain matched scenario. These results highlight that performance differences between domain matched and mismatched models are driven more by neutral classification than by sweep detection.

**Figure 3:**
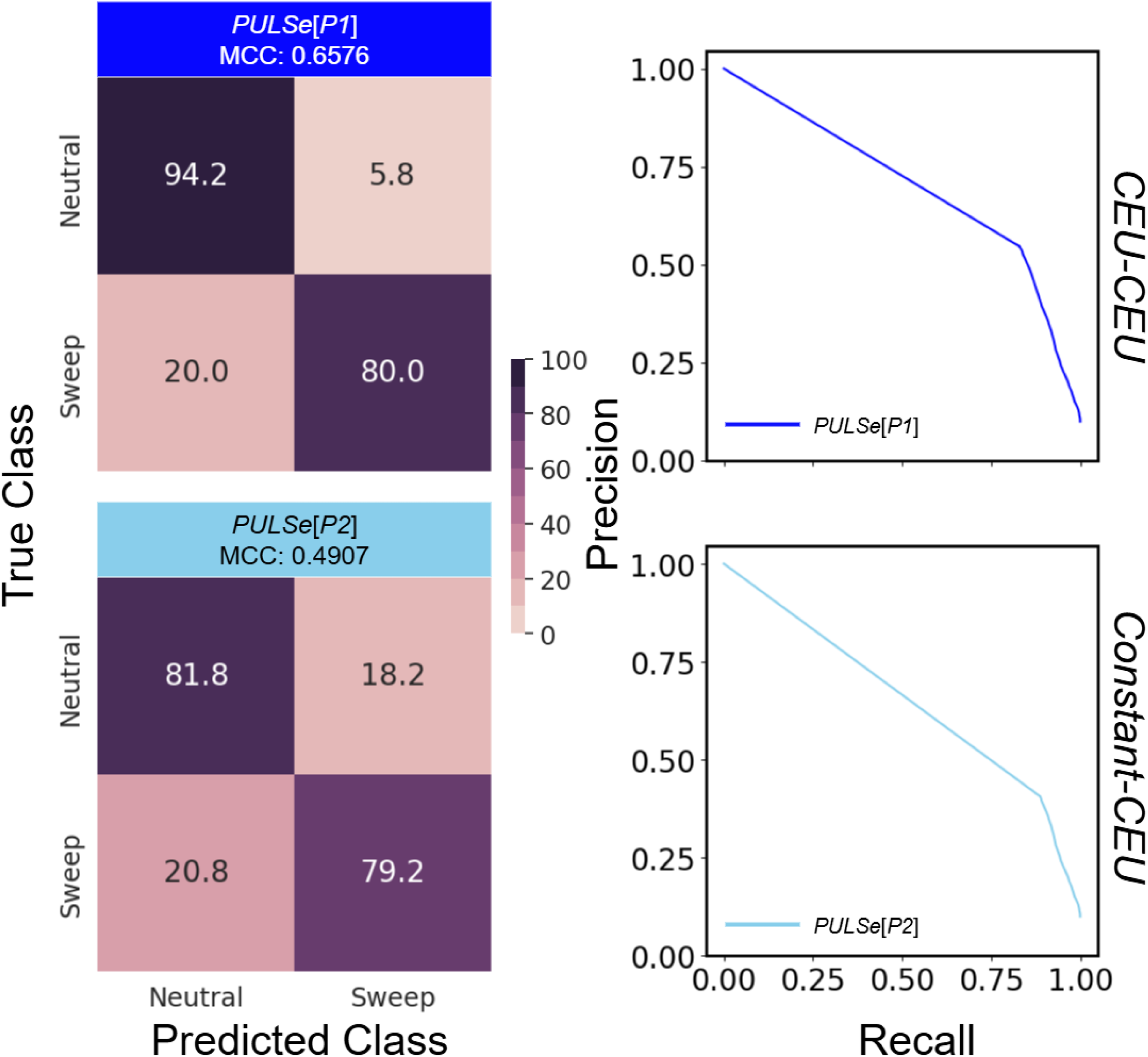
Classification rates and accuracies as depicted by confusion matrices and reliability to detect sweeps as depicted by precision-recall curves to differentiate sweeps from neutrality on the unlabeled set for the *CEU-CEU* (top row) and *Constant-CEU* (bottom row) scenarios (see *Modeling Description* subsection of the *Results*) for the hyperparameter tuned and calibrated *PULSe*[*P1*] and *PULSe*[*P2*] models. For both scenarios, the unlabeled set contains 9,000 neutral and 1,000 sweep replicates simulated based on the recent strong bottleneck demographic history of the CEU human population from the 1000 Genomes Project dataset.

### Genomes also experiencing purifying selection

To evaluate the robustness of *PULSe* when the unlabeled set includes genomic regions affected by other evolutionary forces beyond only positive selection and neutral processes, we incorporated simulated replicates of background selection into the unlabeled pool. Background selection is a process by which purifying selection against deleterious mutations at a set of loci indirectly reduces genetic variation at nearby neutral sites due to linkage [Charlesworth et al., 1993, Hudson and Kaplan, 1995, Charlesworth, 2012]. This setup presents a particularly informative test case, as background selection is a pervasive force [McVicker et al., 2009, Comeron, 2014, Pouyet et al., 2018, Cvijovíc et al., 2018] can produce patterns of genetic variation that can resemble those generated by selective sweeps [Keinan and Reich, 2010, Seger et al., 2010, Nicolaisen and Desai, 2013], thereby potentially complicating binary classification tasks [Huber et al., 2016].

We generated 1,000 background selection replicates using the forward-time simulator SLiM [Haller and Messer, 2019], applying the same genetic and demographic parameters as those used in CEU simulations for training *PULSe*, but with the addition of background selection. Specifically, each 1.1 Mb simulated region contained a 55 kilobase protein-coding gene modeled after the simulated human gene architecture in Cheng et al. [2017], and composed of 50 exons (100 bases each), 49 introns (one kilobase each), 5^′^ and 3^′^ untranslated regions (UTRs; 200 and 800 bases, respectively), chosen to closely match the mean element lengths from the human genome [Mignone et al., 2002, Sakharkar et al., 2004]. Within this gene, recessive (*h* = 0.1) deleterious mutations arose with selection coefficients drawn from a gamma distribution with mean of –0.0294 and shape parameter of 0.184 [Boyko et al., 2008, Schrider and Kern, 2017]. The percentage of mutations that were deleterious in exons, introns and UTRs were 75%, 10%, and 50%, respectively.

Our findings indicate that *PULSe* models under both domain matched and mismatched scenarios are robust toward misclassifying background selection replicates as selective sweeps (Figure 4). This finding is unsurprising given that our image representations are based on haplotype structure, and unlike selective sweeps, background selection typically does not produce the same distinct haplotype patterns [Fagny et al., 2014, Schrider, 2020]. In the domain matched setting, the rate of classifying background selection replicates as non-sweeps (93.4%) nearly mirrors that of the neutral detection rate (93.9%). Interestingly, in the domain mismatched scenario, the rate of classifying background selection replicates as non-sweeps (83.1%) comfortably exceeds the neutral detection rate (79.1%). This result highlights that in the domain mismatched setting, *PULSe* is even less likely to mistakenly classify background selection as a sweep than it is to misclassify neutrally evolving regions, compared to domain matched setting.

**Figure 4:**
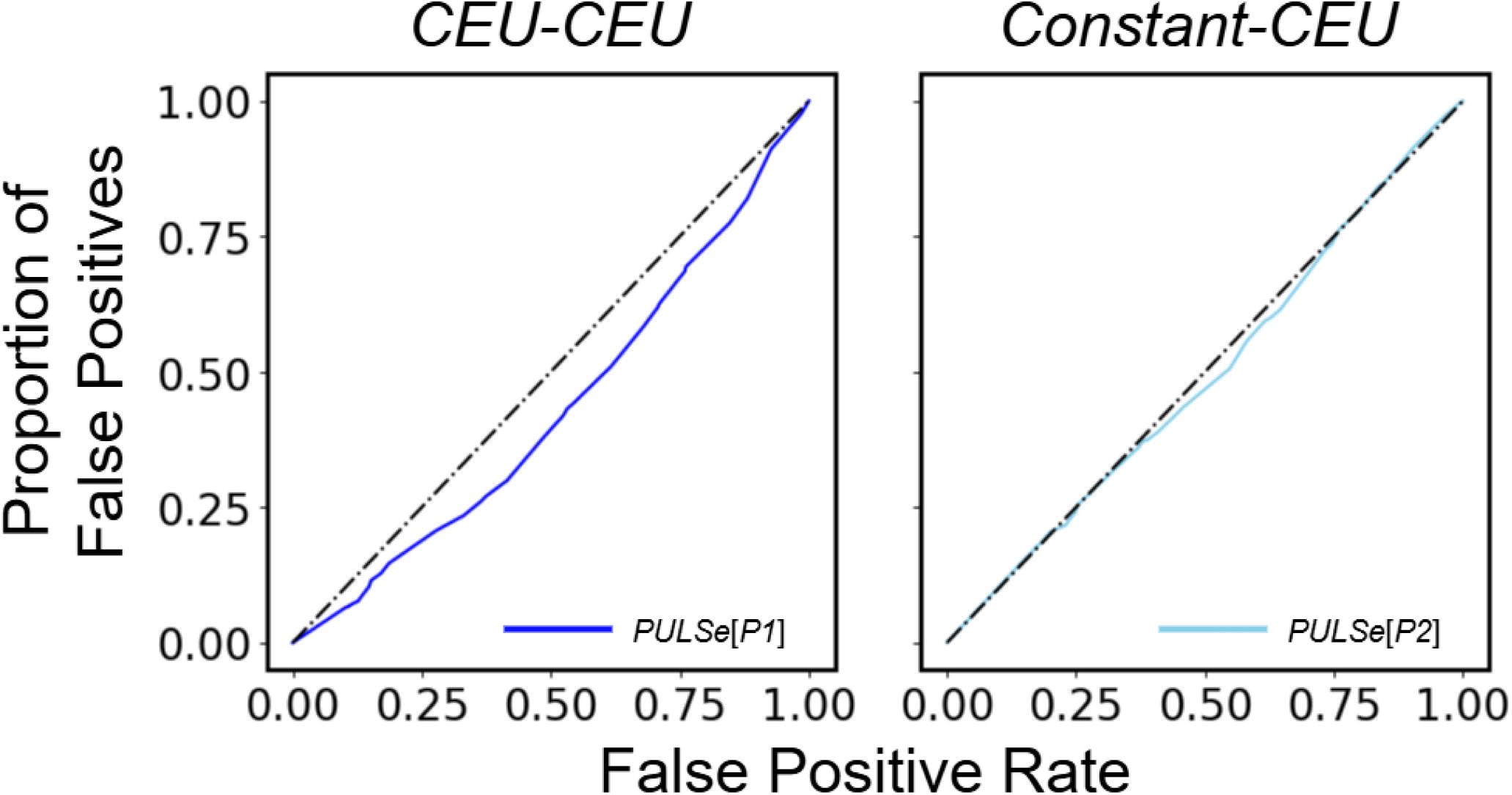
The probability of falsely detecting background selection as a sweep as a function of the false positive rate for the *PULSe*[*P1*] and *PULSe*[*P2*] models applied to the *CEU-CEU* (left) and *Constant-CEU* (right) scenarios, respectively. Trained models are identical to those of Figure 3, with the exception that 1,000 out of 9,000 neutral replicates were replaced by 1,000 background selection replicates. The proportion of false signals is calculated as the fraction of background selection replicates whose sweep probabilities exceed the threshold corresponding to a given false positive rate among neutral replicates.

### Application to unphased data

Though *PULSe* models are originally trained on phased haplotype data, we tested their robustness when trained on unphased multilocus genotype (MLG) data. This is an important practical consideration, especially for nonmodel organisms where phasing is often unreliable or infeasible [Browning and Browning, 2011]. To generate *PULSe* input images from unphased data, we first isolate each pair of consecutive rows, *i.e* haplotypes 2*i* − 1 and 2*i*, where *i* = 1, 2*, …, n/*2, from the original alignments of *n* haplotypes. We then transform each pair of isolated haplotypes into a single unphased MLG by counting the number of minor alleles at each diploid genotype, yielding genotypes coded as 0, 1, or 2. Next, we applied the same transformation and image construction procedures described in the *Image Generation* subsection of the *Materials and Methods* to produce the images, which results in an image of size 99 × 238 for our sample of *n* = 198 haplotypes. We resized these images to 198 × 238, and divided pixel values by a factor of two prior to computing HOG features to avoid altering the feature vector lengths and pixel value ranges used in *PULSe* models. Following image generation and resizing, we computed HOG representations using the *P1* and *P2* pipelines. Using these HOG representations, we trained and hyperparameter tuned (see *Hyperparameter tuning for the base classifier* subsection above for details) two new models, denoted by *PULSe*[*P1*, *MLG*] under the domain matched scenario and *PULSe*[*P2*, *MLG*] under the domain mismatched scenario.

In the domain matched setting, *PULSe*[*P1*, *MLG*] achieved performance nearly identical to its haplotype-based counterpart *PULSe*[*P1*], with approximately 3% lower neutral detection rate and a slightly higher sweep detection rate (compare Figure 5 with Figure 3). Though, its MCC of 0.5904 and AUPRC of 0.6940 were also reduced, the overall results indicate that the model retains strong predictive power on MLG input in the domain match scenario. In contrast, *PULSe*[*P2*, *MLG*] showed a more notable drop under the domain mismatched setting, with a 7% reduction in neutral detection rate, a 5.5% loss in sweep detection rate, and a more pronounced decline in MCC (0.3172) and AUPRC (0.4601) compared to the haplotype setting (compare Figure 5 with Figure 3). Both *PULSe* MLG models were best calibrated with a calibration threshold of 0.3. These results demonstrate that *PULSe* retains meaningful classification performance on unphased data in domain matched settings, though stronger performance when experiencing domain shift may require additional modeling modifications.

**Figure 5:**
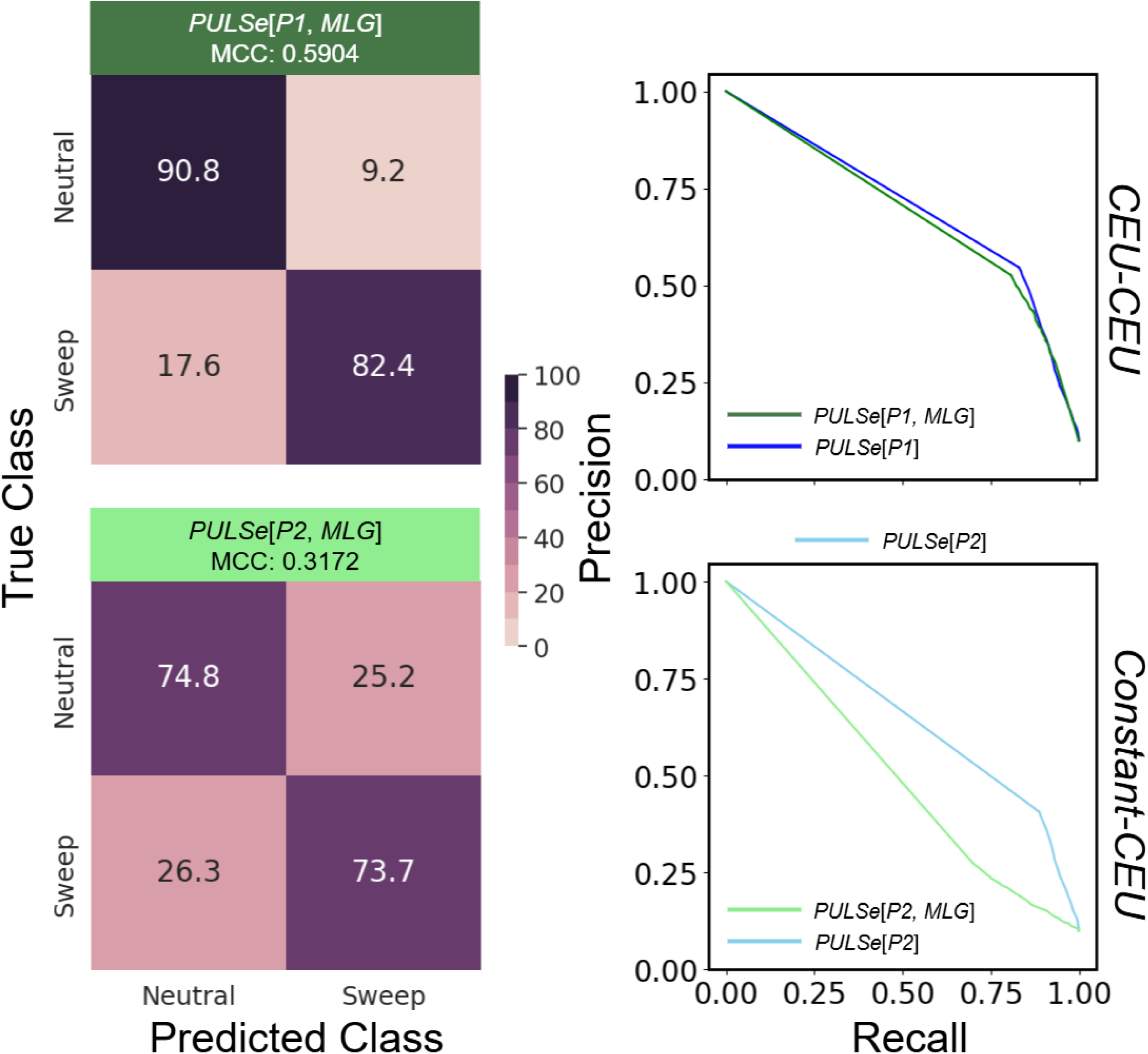
Classification rates and accuracies as depicted by confusion matrices and reliability to detect sweeps as depicted by precision-recall curves to differentiate sweeps from neutrality on the unlabeled set for the *CEU-CEU* (top row) and *Constant-CEU* (bottom row) scenarios (see *Modeling Description* subsection of the *Results*) for the hyperparameter tuned and calibrated *PULSe*[*P1*, *MLG*] and *PULSe*[*P2*, *MLG*] models that operate on variation from unphased multilocus genotypes. For both scenarios, the unlabeled set contains 9,000 neutral and 1,000 sweep replicates simulated based on the recent strong bottleneck demographic history of the CEU human population from the 1000 Genomes Project dataset. The *PULSe*[*P1*, *MLG*] and *PULSe*[*P2*, *MLG*] architectures are identical to that of *PULSe*[*P1*] and *PULSe*[*P2*], respectively, with the difference being that the input is based on unphased multilocus genotype variation instead of phased haplotype variation. The precision-recall curves for the *PULSe*[*P1*] and *PULSe*[*P2*] models are identical to those in Figure 3.

### Effect of lower sample size

Our assessment of *PULSe* models thus far has been performed with a fairly large sample size of 99 diploid individuals. To motivate the understanding of how a comparatively lower sample size affects *PULSe*, we evaluated its performance with roughly a quarter of the sample size of 50 haplotypes instead of 198. We then transformed these alignments into images using the same procedure detailed in the *Image generation* subsection of *Materials and Methods*. The resulting images are of size 50 × 238, which we resized to 198 × 238 prior to computing HOG features to facilitate direct comparison against *PULSe* models trained on 198 haplotypes without introducing confusion regarding different HOG feature vector sizes. Following image generation and resizing, we computed HOG representations using the *P1* and *P2* pipelines. Using these HOG representations, we trained and hyperparameter tuned (see *Hyperparameter tuning for the base classifier* subsection above for details) two new models, denoted by *PULSe*[*P1*, *50h*] under the domain matched scenario and *PULSe*[*P2*, *50h*] under the domain mismatched scenario. This setup allowed us to evaluate the effect of reduced population sample size on classification performance across both training scenarios.

In the domain matched setting, *PULSe*[*P1*, *50h*] exhibited a moderate decline in performance (Figure 6) compared to *PULSe*[*P1*] (Figure 3). The neutral detection rate dropped by approximately 7%, whereas the sweep detection rate improved slightly by over 2% compared to *PULSe*[*P1*]. However, both AUPRC (from 0.727 to 0.6434) and MCC (from 0.6576 to 0.5264) declined, reflecting a loss in overall discriminative ability. Despite this loss, the results remain encouraging, indicating that *PULSe* can still operate effectively with reduced sample sizes under domain matched conditions. In contrast, performance in the domain mismatched setting deteriorated substantially for *PULSe*[*P2*, *50h*]. Both neutral and sweep detection rates fell by over 10%, and AUPRC and MCC respectively dropped by around 19% (0.4630 compared to 0.6531) and 25% (0.2335 compared to 0.4907) relative to *PULSe*[*P2*] (compare Figure 6 with Figure 3). These findings closely mirror those observed for the *PULSe*[*P1*, *MLG*] and *PULSe*[*P2*, *MLG*] models in terms of the sharp decline in domain mismatch performance compared to *PULSe*[*P1*] and *PULSe*[*P2*] models. However, contrary to the findings from the *PULSe*[*P1*, *MLG*] and *PULSe*[*P2*, *MLG*] models, the *PULSe*[*P1*, *50h*] and *PULSe*[*P2*, *50h*] models are best calibrated at a calibration threshold of 0.3.

**Figure 6:**
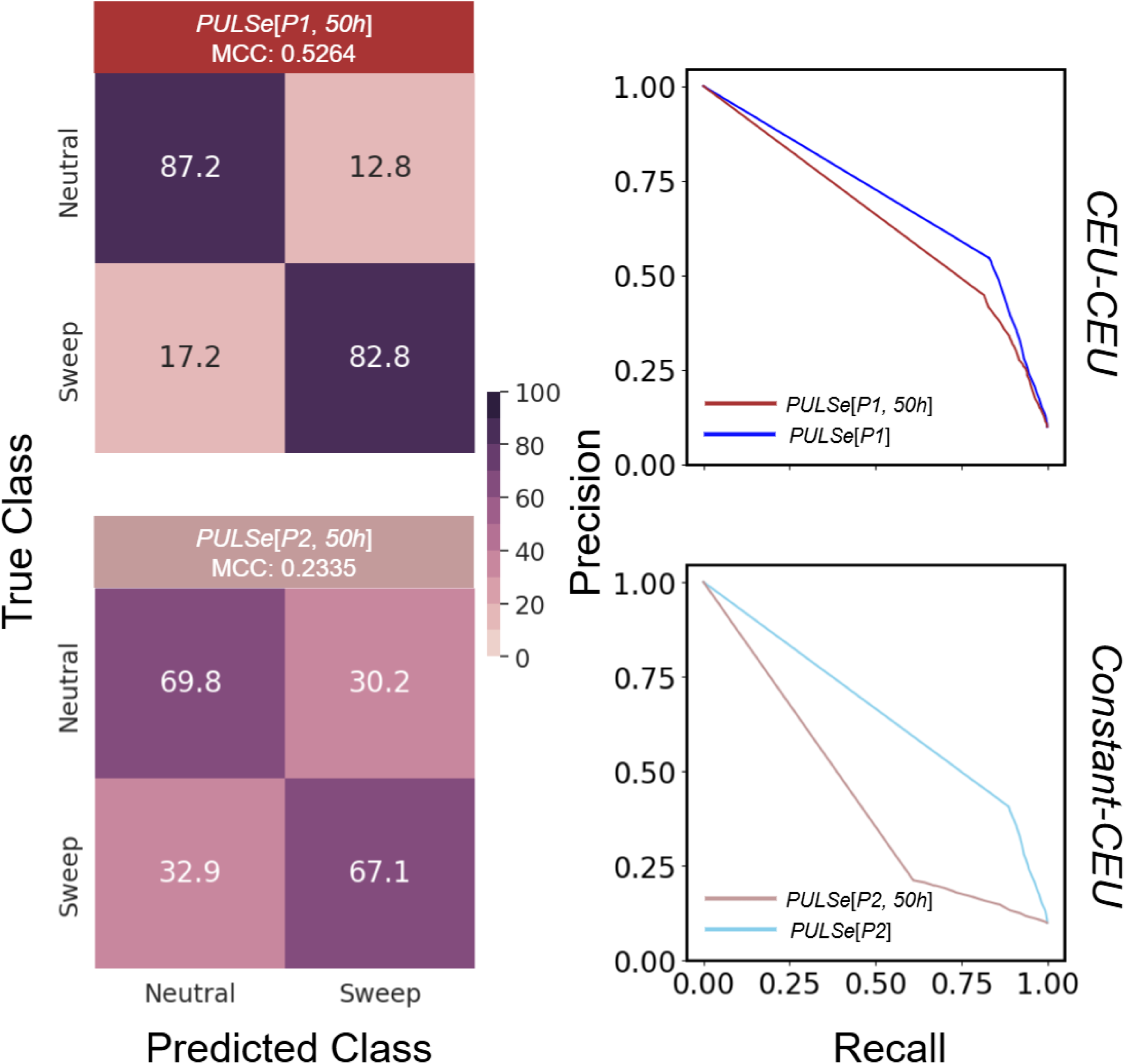
Classification rates and accuracies as depicted by confusion matrices and reliability to detect sweeps as depicted by precision-recall curves to differentiate sweeps from neutrality on the unlabeled set for the *CEU-CEU* (top row) and *Constant-CEU* (bottom row) scenarios (see *Modeling Description* subsection of the *Results*) for the hyperparameter tuned and calibrated *PULSe*[*P1*, *50h*] and *PULSe*[*P2*, *50h*] models that operate on variation from 50 sampled haplotypes. For both scenarios, the unlabeled set contains 9,000 neutral and 1,000 sweep replicates simulated based on the recent strong bottleneck demographic history of the CEU human population from the 1000 Genomes Project dataset. The *PULSe*[*P1*, *50h*] and *PULSe*[*P2*, *50h*] architectures are identical to that of *PULSe*[*P1*] and *PULSe*[*P2*], with the difference being that the input is based on variation from 50 haplotypes instead of 198. The precision-recall curves for the *PULSe*[*P1*] and *PULSe*[*P2*] models are identical to those in Figure 3.

### Benchmarking against alternate architectures

To contextualize the performance of *PULSe*, we benchmarked it against an alternate machine learning architecture under both domain match and mismatch scenarios. In an attempt to benchmark *PULSe* against a traditional domain-adaptive method, we first chose the small multi-branch CNN (*smbCNN*) architecture, which demonstrated strong performance in Arnab et al. [2025]. For context, this architecture is the identical CNN used by Kern and Schrider [2018], except with two classes instead of five and operates on *PULSe* images (see *Image Generation* subsection of the *Materials and Methods*) we developed instead of the native summary statistic images.

Building on this foundation, we explored how gradient reversal layers (GRLs), as implemented in the dadaSIA framework [Mo and Siepel, 2023], can render the *smbCNN* architecture domain adaptive. Mo and Siepel [2023] showed that equipping a previous model (termed SIA) [Hejase et al., 2022] with a GRL substantially improved performance under both domain matched and mismatched conditions. Prior work in domain adversarial learning has shown that GRLs allow neural networks to learn features that are both predictive of labels in the source domain and invariant to differences between source (training) and target (testing) domains [Ganin et al., 2016, Ding et al., 2019, Zügner et al., 2020]. By inserting a GRL-connected domain discriminator into the *smbCNN* architecture, we adapted the network to jointly minimize classification loss on labeled source samples and maximize domain classification loss across both source and target samples, encouraging the feature extractor to obtain domain-invariant representations. This modification may allow the original architecture to function robustly across mismatched domain conditions without explicitly modeling population demographic history. Notably, we chose not to compare *PULSe* directly against dadaSIA, as it operates on ancestral recombination graphs (ARGs) as input that are fundamentally different from the *PULSe* images, thereby avoiding confounding differences arising from input types such as summary statistics or ARG features.

We denote the GRL-equipped alternate *smbCNN* architecture that takes *PULSe* images (*PI*) as input by *smbCNN GRL*[*PI*]. A direct comparison with *PULSe* is not possible as *smbCNN GRL*[*PI*] requires labeled samples from both classes for training, whereas *PULSe* operates with the labeled positive sweep class only. Nevertheless, this test provides a useful performance baseline even though *smbCNN GRL*[*PI*] benefits from a significant advantage in training supervision. A detailed descrip-tion of the *smbCNN GRL*[*PI*] model architecture and training procedure is provided in the *Training the smbCNN GRL[PI] architecture* subsection of the *Materials and Methods*.

In the domain matched scenario, *smbCNN GRL*[*PI*] outperformed *PULSe*[*P1*] across all evaluation metrics. It achieved a neutral detection accuracy of 93.3%, sweep detection accuracy of 88.2%, an AUPRC of 0.833, and an MCC of 0.6917 (Figure 7), all of which exceeded the corresponding metrics for *PULSe*[*P1*] (Figure 3). However, in the domain mismatch setting, its performance degraded substantially relative to *PULSe*[*P2*]. Neutral detection accuracy dropped dramatically to 67.3%, sweep detection rose slightly to 89.5%, with a lower AUPRC of 0.577 (compared to 0.6531 of *PULSe*[*P2*]) and a lower MCC of 0.3505 (compared to 0.4907 of *PULSe*[*P2*]) (compare Figure 7 with Figure 3), all of which except for sweep detection accuracy were markedly lower than those achieved by *PULSe*[*P2*] (Figure 3). Notably, the domain output of *smbCNN GRL*[*PI*] identified 55.95% of samples as out-of-domain under domain mismatch with predicted domain probability higher than 0.5, whereas indicating 0% out-of-domain with all samples having predicted domain probability lower than 0.5, under domain match (Figure S7), underscoring its capacity to detect domain shifts. However, despite recognizing this domain shift, the model was evidently not sufficiently effective at classifying samples in the alternate domain compared to *PULSe*.

**Figure 7:**
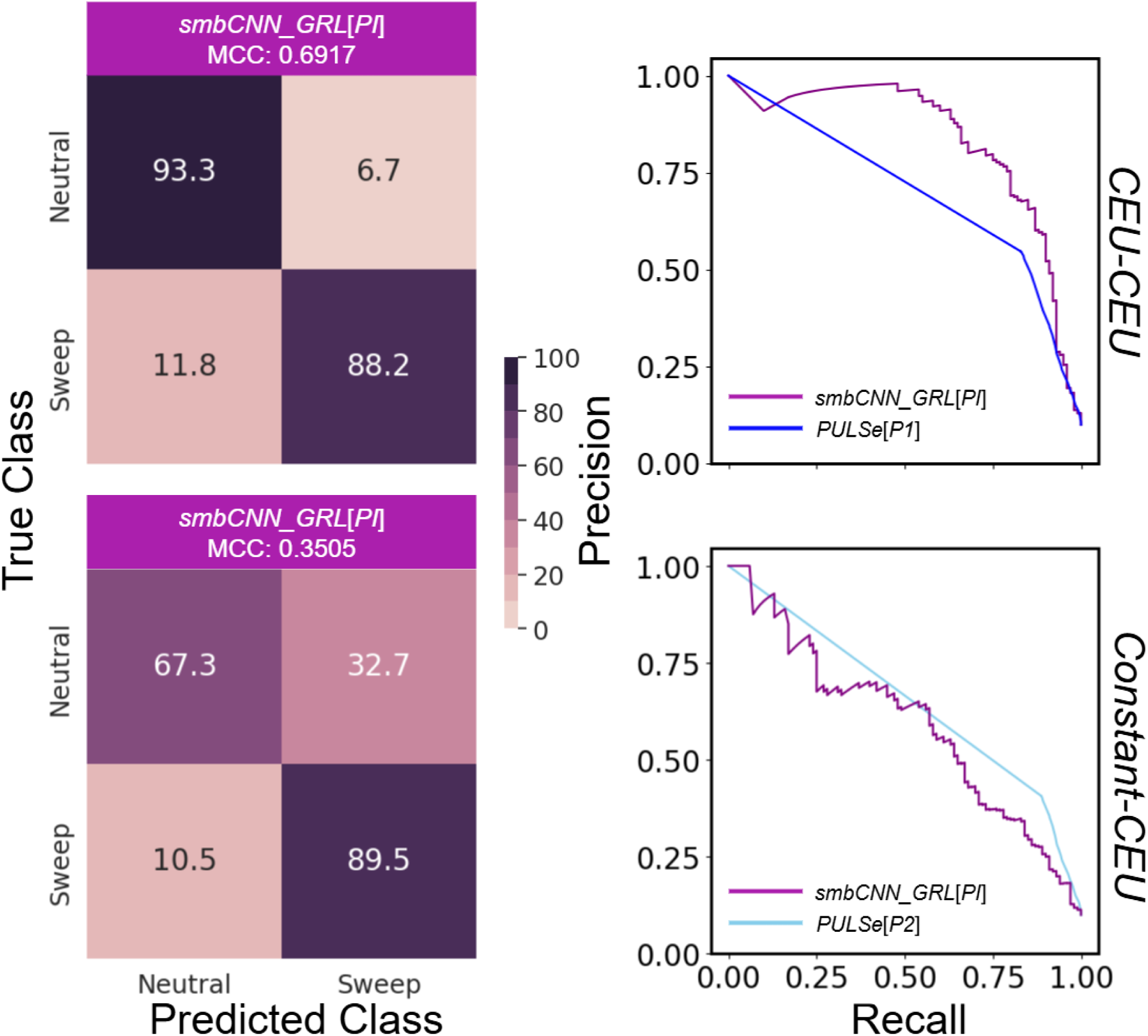
Classification rates and accuracies as depicted by confusion matrices and reliability to detect sweeps as depicted by precision-recall curves to differentiate sweeps from neutrality on the unlabeled set for the *CEU-CEU* (top row) and *Constant-CEU* (bottom row) scenarios (see *Modeling Description* subsection of the *Results*) for the *PULSe*[*P1*] and *PULSe*[*P2*] models, compared against *smbCNN GRL*[*PI*] (see *Benchmarking against alternate architectures* subsection of the *Results* for details). For both scenarios, the unlabeled set (test set in the case of *smbCNN GRL*[*PI*] models) contains 9,000 neutral and 1,000 sweep replicates simulated based on the recent strong bottleneck demographic history of the CEU human population from the 1000 Genomes Project dataset. The precision-recall curves for the *PULSe* models are identical to those in Figure 3.

### Application to human genomes of known European demographic history

To explore empirical signals of adaptation, we applied *PULSe* to phased haplotype data from 99 individuals in the CEU population from the 1000 Genomes Project dataset [The 1000 Genomes Project Consortium, 2015]. As detailed in the *Empirical data filtration and image generation* sub-section of the *Materials and Methods*, we first excluded low-mappability regions and partitioned each chromosome into overlapping 499-SNP windows with a stride of 50 SNPs. Each window was rendered as an image of spatial haplotype variation, and HOG features were extracted using both domain matched (*P1*) and domain mismatched (*P2*) pipelines. These windows were then used as the unlabeled set to the calibrated logistic regression models of *PULSe*[*P1*] and *PULSe*[*P2*].

Using the *P1* pipeline, 3.94% of windows exceeded the sweep probability threshold of 0.5, with 3.42% of windows assigned a probability of exactly 1.0. In contrast, the *P2* pipeline, trained under a mismatched demographic model, classified 8.57% of windows above the 0.5 threshold, and 8.52% received a predicted probability of 1.0. This inflation of high-confidence predictions under domain mismatch reflects the impact of the calibration methodology that we applied. In total, the high sweep probability windows with predicted sweep probability exceeding 0.5 found by *PULSe*[*P1*] fall within the bounds of 954 genes in our genome-wide scan, whereas the peaks identified by *PULSe*[*P2*] fall within the bounds of 1,834 genes.

Of the genes identified by *PULSe*[*P1*] and *PULSe*[*P2*], a total of 609 genes, which is around 64% of the *PULSe*[*P1*] discoveries, are shared between the two scans. The fact that *PULSe*[*P2*], despite being handicapped by domain mismatch, still recovered the majority of discoveries of *PULSe*[*P1*] reinforces confidence in the reliability of the *PULSe*[*P2*] results. The CEU scan thus provides a valuable opportunity to assess the utility of *PULSe*[*P2*] in a true domain mismatched application, as the CEU population has been extensively studied [Szpiech and Hernandez, 2014, Duforet-Frebourg et al., 2016, Chen et al., 2016, Kern and Schrider, 2018, Mughal et al., 2020, Wang et al., 2021, Arnab et al., 2025], and offers a strong comparative basis for evaluating our findings. Among the *PULSe*[*P2*] identified candidates are several well-established targets of positive selection in European populations, including *LCT*, which underlies lactase persistence and has long been recognized as a hallmark of recent adaptation to dairy farming [Sabeti et al., 2006, Bersaglieri et al., 2004]. The finding of this signal lends credence to the ability of *PULSe* to recapitulate known adaptive loci. Another notable candidate is *ABCA12*, a gene essential to skin barrier function through its role in lipid transport in the epidermis [Akiyama, 2014, Colonna et al., 2014]. Adaptation at this locus has been linked to variation in environmental ultraviolet exposure and skin physiology, particularly among Eurasian populations. *PULSe*[*P2*] also detected selection at *HLA-DRB6*, a class II pseudogene located in the major histocompatibility complex (MHC) region, where long-term balancing and intermittent positive selection events are hypothesized to maintain diversity for pathogen defense [Cree et al., 2010]. Though *HLA-DRB6* is nonfunctional, its genomic context suggests it may be linked to selective pressures acting on nearby regulatory or coding regions within the MHC.

It is important to note that all of the above candidates were also found by the *PULSe*[*P1*] scan. Interestingly, the common gene set also includes previously identified, but less well-characterized, candidates such as *RBMS3* and *NKAIN2*, both of which were also highlighted in earlier scans by the *TrIdent* classifier of Arnab et al. [2025]. This convergence is particularly notable given that *PULSe* and *TrIdent* operate on almost identical image representations of haplotype variation, differing primarily in their downstream modeling architectures. *RBMS3* has been implicated as a tumor suppressor in breast cancer and may play roles in transcriptional regulation and cell cycle arrest [Yang et al., 2018], whereas *NKAIN2* has been linked to tumor inhibition and neurological function [Zhao et al., 2015]. The recovery of these cancer-associated genes from independent scans suggests the possibility that some adaptive signals may be linked to pleiotropic effects involving disease resistance or cell proliferation. Taken together, these overlapping results reinforce the capacity of *PULSe* to uncover both canonical and novel candidates of positive selection, and offer a biologically meaningful set of targets for downstream functional and evolutionary investigation.

This consistency across pipelines further underscores the ability of *PULSe* to recover biologically meaningful candidates of positive selection and reinforces its ability to identify our targeted mode of selection across modeling conditions. Notably, over half of the genes identified by *PULSe*[*P1*] were also recovered by *PULSe*[*P2*], demonstrating a degree of robustness despite demographic model misalignment. The increased number of candidate sweeps detected in *PULSe*[*P2*] reflects a higher false positive rate, which is expected given that the labeled sweeps from the constant-sized demographic history lack the recent population size bottleneck present in the CEU population.

To investigate whether identified sweep candidates were enriched for particular biological functions or structures, we conducted Gene Ontology (GO) enrichment analysis using GOrilla [Eden et al., 2009] on two unranked lists of genes, where the target list contains candidate genes detected by the domain mismatched scenario of *PULSe*[*P2*], and the background list includes all genes that are not removed by our filtration process. Significant enrichment was determined using an false-discovery rate-adjusted *q*-value threshold of 0.05. Among the 240 significant biological process terms identified (Table S1), several top-ranked categories point to broad mechanisms of cellular structure management, such as “cellular component organization”,“cellular component organization or biogenesis”, and “regulation of cell morphogenesis”. These terms may suggest selection on genes influencing the organization and regulation of physical cell structures. The 95 enriched molecular function terms (Table S2) highlighted key activities such as “ion binding”, “cytoskeletal protein binding”, and “GTPase binding”, all of which play essential roles in signaling and maintaining cellular integrity. Meanwhile, enriched cellular component terms (Table S3) emphasized structures like “neuron part”, “cell projection”, and “plasma membrane bounded cell projection part”, reinforcing the notion that many identified sweep candidates may affect cell morphology and intercellular communication. The GO analyses highlight a broad but consistent enrichment of genes involved in cellular structure organization, signaling, and membrane-associated components, suggesting that positive selection in the CEU population may have acted on pathways essential for maintaining and regulating cell architecture and communication.

### Application to human genomes of unknown South Asian demography

To extend our evaluation of *PULSe* beyond the CEU population, we conducted a genome-wide scan of the Bengali in Bangladesh (BEB) population containing phased haplotype data from 86 individuals from the 1000 Genomes Project dataset [The 1000 Genomes Project Consortium, 2015] using the same domain mismatched pipeline *PULSe*[*P2*] that was previously applied to the CEU. This test case followed the identical training scheme and calibration approach used in the CEU scan.

The populations within Bangladesh exhibit high levels of genetic variation, characterized by contributions from a broad array of ancestral groups [Reich et al., 2009, Gazi et al., 2013, Narasimhan et al., 2019]. These contributions include overlapping ancestries from South and East Asia, the Middle East, and Europe, which together contribute to a complex admixture landscape. This ancestry mixture has been shaped by centuries of trade, invasions, warfare, colonization, and religious migration, all of which introduced novel ancestral lineages into the region [Reich et al., 2009, Prakash, 2014, Basu et al., 2016, Narasimhan et al., 2019]. Despite this heterogeneity, the population shows minimal substructure that would reflect isolated population clusters [Chatterjee, 2020]. Instead, ancestral components are distributed broadly and continuously across the population, reflecting a long history of migration, admixture, and demographic change. Additionally, the population has experienced repeated natural disasters and recurrent exposure to infectious diseases, both of which may have exerted further selective and demographic pressures [Sen, 1982, Haque, 1997, Karlsson et al., 2013]. Such demographic complexity makes a BEB genomic dataset an excellent testbed for assessing robustness of *PULSe* under domain mismatch, especially given admixture and history of environmental and sociopolitical disruptions experienced by the population.

*PULSe* identified 9.77% of windows exceeding the sweep probability threshold of 0.5, with a majority of those windows receiving a predicted probability of 1.0. This elevated detection rate compared to the CEU scan likely reflects the greater genetic variation of the BEB population, shaped by its complex demographic history, which makes even weaker sweeps stand out more prominently in the genomic landscape. Overall, the identified sweep probability peaks intersect the boundaries of 1,978 genes.

Among the most compelling sweep candidates detected are *KCNH3*, *KCNH5*, and *KCNH7* on chromosomes 12, 14, and 2, respectively, all of which encode voltage-gated potassium channels that regulate the flow of potassium ions across cell membranes [Jensen et al., 2012]. These channels are essential for maintaining neuronal excitability and proper synaptic transmission, playing a critical role in electrical signaling within the brain [Singh and Auerbach, 2024]. Their involvement in neurodevelopmental processes is well established, and they have been implicated in susceptibility to various cognitive and psychiatric disorders, including schizophrenia, bipolar disorder, and dementia [Strauss et al., 2014, Rexach et al., 2023, Wang et al., 2019]. *KCNH3*, in particular, has previously been identified as a shared sweep candidate between closely related South and East Asian populations [Liu et al., 2017]. This finding is especially notable in light of recent studies reporting increased prevalence of certain mental health disorders among South Asian populations [Vidyasagaran et al., 2023, Islam et al., 2024]. Just as our CEU scan identified cancer associated genes, we hypothesize that certain adaptations in South Asians, initially advantageous under past environmental stressors, may now predispose individuals to deleterious conditions in modern environments. In addition to their neurological roles, *KCNH5* and *KCNH7* have also been linked to susceptibility to severe cholera in the BEB population [Karlsson et al., 2013], suggesting a possible dual role in adaptation that balances neurological function and resistance to infectious diseases in the pathogen-rich environment of Bangladesh.

Situated on chromosome 20, *DOK5* (Docking Protein 5) is another sweep candidate identified by *PULSe* that plays a role in insulin and neurotrophin signaling pathways [Cai et al., 2003]. *DOK5* has been associated with type II diabetes and metabolic disorders, and has been identified as a candidate for positive selection in a scan of Indian populations [Metspalu et al., 2011], who are also from South Asia. Moreover, located on chromosome 15, *SLC24A5* emerged as another strong candidate signal of selection in our scan. This gene encodes a cation exchanger involved in melanosome function and is one of the most robustly documented targets of positive selection in human populations outside of Africa [Lamason et al., 2005]. An allele at locus rs1426654 of *SLC24A5* that contributes to lighter skin pigmentation is at intermediate to high frequencies in South Asia [Mallick et al., 2013]. Furthermore, *PULSe* also detected *OAS1* on chromosome 12 as a potential candidate, a gene that plays a central role in the innate immune response. The antiviral mechanism of *OAS1* [Melchjorsen et al., 2009] is critical for controlling infections such as dengue, hepatitis, and other RNA viruses, many of which are endemic to South Asia [Kristiansen et al., 2010]. *OAS1* is also a hypothesized target of selection in primates [Mozzi et al., 2015], and in humans it is thought to have been introgressed from Neanderthals [Sams et al., 2016]. *PULSe* also detected multiple candidate genes within the MHC region, including *HLA-DRB1*, *HLA-DRB5*, and *HLA-DQB1*. The MHC region is a well-established hotspot of selection in human populations due to its central role in host-pathogen coevolution and pathogen defense [Trowsdale, 2011]. The identification of sweep signals in this region within BEB further supports the involvement of this region in adaptation to local pathogenic environments.

Our BEB scan also revealed sweep candidates that overlapped with those identified in the CEU population with the *P2* pipeline, including *LCT*, *ABCA12*, *RBMS3*, and *NKAIN2*. The discovery of these genes across populations with distinct evolutionary histories suggests that they may represent robust targets of positive selection, either due to convergent pressures or shared ancestral adaptations. In particular, the replication of candidates like *RBMS3* and *NKAIN2*, indicates that certain adaptive signals may persist across different environments. These shared findings bolster the hypothesis that some adaptive variants may have broad functional relevance, potentially tied to disease resistance.

Because we did not conduct simulated benchmarking involving a BEB demographic model, including sample size, the efficacy of the calibration method employed was not directly validated for this test case. Instead, we applied the same probability threshold (0.3) that was optimized for the *Constant-CEU* scenario. Our testing on the CEU demographic history (see *Improving detection performance with model calibration* subsection above) indicated that increasing the threshold reduces false positives but may reduce sensitivity to detect true sweeps. To assess the robustness of our BEB scan under alternative calibration parameters, we tested thresholds of 0.4 and 0.5. With a threshold of 0.4, the number of genes with predicted sweep signals decreased to 1,124, and with a cutoff of 0.5, the total declined further to 556. Notably, even under this strictest setting, only *LCT*, *ABCA12* and *SLC24A5* were excluded, suggesting that these were likely weaker signals within the BEB scan.

To better understand the biological context of sweep candidates in the BEB, we carried out GO enrichment analysis using GOrilla [Eden et al., 2009]. Candidate genes identified by *PULSe* were compared against a background set of all retained genes following our filtration procedure, with significance defined by an FDR-adjusted *q*-value threshold of 0.05. We did not identify any notable overlaps between the CEU and BEB GO terms. Among the 289 significantly enriched biological process terms (Table S4), several of the top categories reflected processes involved in structural organization and neuronal development, including “cellular component organization”, “regulation of plasma membrane bounded cell projection organization”, and “regulation of neuron projection development”. These findings suggest that some candidate sweep signals may indicate potential selection on genes influencing neural connectivity or cell morphology. The 95 enriched molecular function terms (Table S5) were heavily weighted toward general binding activities, such as “ion binding”, “protein binding”, and “anion binding”. In terms of cellular component enrichment (Table S6), top terms such as “neuron part”, “cell projection part”, and “membrane” indicate a strong representation of genes associated with cellular extensions and membrane-associated functions. These results suggest that many of the BEB sweep candidates may play roles in shaping how cells interact with their environment, particularly in the context of neural and structural function.

## Discussion

*PULSe* exhibited strong performance in the test case of selective sweep detection, which served as a proof of concept for evaluating the framework. In the domain matched setting, similar to the single set training protocol described by Elkan and Noto [2008], the demographic history of the training samples mirrored that of the test data. Under these conditions, *PULSe* exhibited comparable performance to the domain adaptive approach of *smbCNN GRL*[*PI*], even though *PULSe* was trained solely with data from the target class. When extended to the domain mismatched scenario, analogous to the case-control design in Elkan and Noto [2008], *PULSe* interestingly outperformed this domain adaptive method. This advantage emerged even under substantial demographic shifts between training and test samples, despite PU learning not being explicitly developed to address domain shift.

When comparing *PULSe* against *smbCNN GRL*[*PI*], our intention was not to position them in a direct head-to-head contest. Such a comparison would be inherently unfair, as training *sm-bCNN GRL*[*PI*] leverages explicit information about the background class through labeled neutral replicates, which is an advantage absent in *PULSe* that operates without any labeled background data. Instead, our primary goal was to examine how each approach responds to domain shifts by comparing their performance changes between domain matched and mismatched settings. Our initial experiments showed that *smbCNN GRL*[*PI*] outperformed the same architecture without GRLs, consistent with the findings of Mo and Siepel [2023]. However, we found that the perfor-mance deterioration of *smbCNN GRL*[*PI*] under domain mismatch, reflected by sharper declines in AUPRC, MCC, and class-specific detection rates, surpassed the relative declines observed for *PULSe*. Several factors might underlie this pronounced performance drop. First, a key reason may lie in how domain adversarial networks function, as they aim to align the marginal distributions of the source and target feature representations globally, effectively forcing the feature extractor to produce domain-invariant representations. This strategy works well when the domain shift is expected to be relatively uniform across classes [Ganin et al., 2016]. However, the interplay between natural selection and demographic history means that domain shifts may often manifest differently for non-neutral and neutral loci. Demographic bottlenecks or expansions can accentuate or obscure non-neutral regions in ways that do not uniformly affect all genomic regions, often altering patterns of variation and linkage disequilibrium that resemble the signatures of selection [Johri et al., 2022]. As a result, regions that are truly neutral in the target domain may still display features that look like sweeps under the feature representations learned from the source domain training data. Because domain adversarial networks primarily align the overall feature distributions without explicitly disentangling class-specific shifts, this may cause the model to confuse these neutral regions for sweeps. This drawback likely contributed to *smbCNN GRL*[*PI*] disproportionately misclassifying many neutral samples under the domain mismatched scenario.

Given the complexities of natural populations, the unlabeled dataset we used in this study intentionally featured a strong imbalance, with the majority of samples representing non-sweep regions. This imbalance is supported by several empirical scans, which suggest that loci evolving under positive selection leading to sweeps are relatively rare [Hernandez et al., 2011, Lohmueller et al., 2011, Granka et al., 2012, Jensen et al., 2019]. Due to the significant class imbalance in our unlabaled sets, to select the optimal *PULSe* architecture and to compare its performance against *smbCNN GRL*[*PI*], we prioritized AUPRC as our primary evaluation metric, instead of the more commonly used receiver operating characteristic (ROC) curve and its area under the curve (AUROC). While ROC curves depict true positive rate as a function of false positive rate to provide a view of performance across all classification thresholds, they treat the positive and negative classes equally [Carter et al., 2016, Chicco and Jurman, 2023]. As a result, ROC curves often fail to adequately reflect performance in situations with significant class imbalance [Chicco and Jurman, 2023]. In contrast, precision-recall curves specifically characterize how well a classifier detects the positive class depicting precision (positive predictive value) against recall (true positive rate), and the AUPRC explicitly summarizes the trade-off between these two measures [Ozenne et al., 2015, Hancock et al., 2023]. This characteristic makes precision-recall curves particularly sensitive to the detection of the positive class [Ozenne et al., 2015, Hancock et al., 2023].

ROC curves for *smbCNN GRL*[*PI*] on the unlabeled set for both domain matched and mis-matched scenarios showcase higher power across most false positive rates compared to that of *PULSe* (Figure S8). The AUROC score comparison contradicts that of AUPRC scores, as in the domain mismatched scenario *PULSe* clearly outperforms *smbCNN GRL*[*PI*] in terms of AUPRC. The conflicting AUPRC and AUROC scores on the domain mismatched setting highlight the drawback if we had used AUROC as the primary performance metric to evaluate *PULSe*. Here, AUROC paints an overly optimistic picture of the performance of *smbCNN GRL*[*PI*], overlooking its failure to detect the neutral (majority) class. However, in *PULSe* applications in which we may expect reasonable balance between the target and background classes, the AUROC should be used as the primary metric for model selection and evaluation.

To extract useful features for input to *PULSe*, we employed HOG [Freeman and Roth, 1995], a legacy computer vision technique that encodes spatial autocovariance by capturing local gradients and edge directions, effectively preserving spatial relationships of pixel values in *PULSe* images that simple image flattening would not be able to preserve [Elkan and Noto, 2008]. Moreover, HOG produces feature arrays of fixed size for a given image dimension, a property that many contemporary approaches do not guarantee without additional pooling or resizing [Lowe, 2004b, Bay et al., 2006, Rublee et al., 2011]. Modern alternatives such as SIFT (Scale Invariant Feature Transfrom; [Lowe, 2004b]), SURF (Speeded-Up Robust Features; [Bay et al., 2006]), or ORB (Oriented FAST and Rotated BRIEF; [Rublee et al., 2011]) offer robust scale and rotation invariance. However, these benefits come at a higher computational cost and are unnecessary for *PULSe* images, where neither scale nor orientation is expected to greatly vary. On the other hand, if genomic variation is summarized into one-dimensional vectors, such as summary statistics, then we do not anticipate any need for further transformation to fit the base classifier of *PULSe*. Conversely, if working with multi-dimensional arrays, then users are not restricted to using HOG. Any feature extraction method can be employed, provided it yields a consistent feature length across samples and ensures that each feature position represents the same characteristic for all samples.

To showcase its practical utility and flexibility, we applied *PULSe* to two distinct empirical contexts. We first focused on the well-studied CEU population, which served as a baseline to understand how *PULSe* behaves when deployed through its two pipelines. Using *PULSe*[*P1*], with labeled positives derived from sweep replicates simulated under an inferred CEU demographic model, we successfully recovered many well-established selective sweep candidates. Meanwhile, *PULSe*[*P2*], trained on sweep replicates generated from a constant population size demographic model, was still able to recapitulate a substantial portion of the sweep candidates identified by *PULSe*[*P1*]. Even with the introduction of the signature population bottleneck of the CEU population, *PULSe*[*P2*] identified the majority of literature-supported candidates found by *PULSe*[*P1*], albeit with an increased number of sweep discoveries. This high degree of overlap in their candidate genes provides support for the robustness of the assessed pipelines, lending confidence that the two *PULSe* pipelines do not fundamentally alter the empirical signals detected. Furthermore, it is worth emphasizing that these analyses were conducted using calibration thresholds optimized for the best calibrated state of each pipeline. However, calibration thresholds can be adjusted to identify only the strongest candidates, or to cast a wider net with less strict cutoffs.

Our application of *PULSe* to the BEB population was potentially the more challenging, yet more significant of the two test cases, particularly because of its intricate demographic history. Positioned between South and Southeast Asia, Bengal has experienced complex layers of ancestral contributions, from early Austroasiatic and Dravidian settlers [Basu et al., 2016, Silva et al., 2017] to moderate Indo-Aryan migration [Metspalu et al., 2011, Narasimhan et al., 2019], centuries of rule by Persian, Turkic, and Central Asian dynasties [Eaton, 1993], and more recent European colonial contact [Prakash, 2014, Chatterjee, 2020]. These historical movements were further compounded by distinct gene flow events that left Bengali individuals with particularly high proportions of East Asian ancestry among South Asians [Reich et al., 2009, Gazi et al., 2013]. The population has also endured repeated, though relatively more recent demographic upheavals through famines, cyclones, and seasonal floods [Sen, 1982, Haque, 1997, Mirza, 2002], massive displacements during the partition of India, and the 1971 Bangladesh Liberation War [Chatterji, 2007, Ranjan, 2016], all of which likely induced bottlenecks, founder effects, and admixture signals that complicate genetic landscapes. Despite this rich and challenging demographic backdrop, and the fact that they represent a large fraction of the global population, the Bengali people remain underexplored in population genomics. By applying *PULSe* in this context, we showcase its ability to operate in complex populations, providing insights that could inform more targeted studies on adaptation and population history in this and similar groups.

Fundamentally, it is of high importance to place *PULSe*, and more broadly, PU learning, within its practical application space, highlighting both its strengths and limitations. In our domain matched experiments, the supervised *smbCNN GRL*[*PI*] model outperformed *PULSe*, as expected given its explicit use of labeled information about the neutral (negative) class. However, *PULSe* required no such negative class data, relying solely on samples of the class of interest, which substantially simplifies study design and minimizes assumptions about the demographic or selective forces shaping the rest of the genome. This advantage makes *PULSe* particularly appealing when simulating or defining realistic negative classes is infeasible. Its straightforward design also means that predictions are easier to interpret, especially when the negative class fails to encompass all evolutionary and non-evolutionary dynamics expected to impact genetic variability in a population. However, most notably, our results indicate that *PULSe* is particularly helpful under domain mismatch scenarios—settings where demographic shifts or sampling differences might undermine more traditional supervised or domain-adaptive approaches. Here, the ability to adjust the calibration threshold becomes invaluable, lending control over the balance between stringency and flexibility, from only capturing the strongest candidates of the target class to broadly flagging all regions suggestive of positive selection. Taken together, these characteristics underscore the versatility of *PULSe* and its promise for evolutionary inference across an array of study systems. Looking forward, an intriguing avenue for future exploration is whether rather than relying exclusively on simulations for the labeled set, one could consider generating augmented copies of well-characterized adaptive regions from empirical genomes of the analyzed population. Such an approach would effectively place the analysis in a domain matched setting, as the empirical images provide direct examples from the expected positive class distribution, which is a scenario where *PULSe* achieves its highest power.

## Materials and Methods

### Simulation protocol

We employed the coalescent simulator discoal [Kern and Schrider, 2016] to generate both neutral and sweep replicates under two demographic models: a nonequilibrium demographic history estimated for European (CEU) humans [Tennessen et al., 2012] and a constant population size model (*Constant*). The CEU demographic model incorporates a recent severe population bottleneck, capturing the complex evolutionary dynamics of this population, whereas the *Constant* model permits investigation of selection under a stable population size assumption.

To simulate selective sweeps, we considered per-generation selection coefficients (*s*) drawn uni-formly at random within the range [0.005, 0.1], initial beneficial allele frequencies (*f*) sampled uni-formly at random within [1*/*(2*N_e_*), 0.2], and fixation times (*τ*) uniformly distributed within [0, 2,000] generations before sampling [Schrider and Kern, 2017]. These choices ensure that our simulations capture a broad array of sweep dynamics, including varying levels of selection strength, hard versus soft sweeps, and different temporal stages of fixation, which all contribute to the detectability of selective events in genetic data.

Both neutral and sweep replicates were simulated with per-site per-generation mutation rates (*µ*) sampled uniformly at random within the interval [2.21×10^−9^, 2.21×10^−8^], with a mean of 1.21×10^−8^ [Scally and Durbin, 2012, Schrider and Kern, 2017]. The per-site per-generation recombination rate (*r*) was drawn from an exponential distribution with a mean of 10^−8^ and truncated at three times the mean [Payseur and Nachman, 2000, Schrider and Kern, 2017]. These settings ensure that our simulations incorporate realistic levels of mutation and recombination heterogeneity.

For each replicate simulated under the inferred CEU demographic history, we sampled 198 haplotypes of length 1.1 megabases (Mb), reflecting the empirical dataset structure. Similarly, for the *Constant* demographic model, we maintained the same simulation parameters and number of haplotypes, except that the effective population size was drawn uniformly at random from *N_e_* ∈ {3,000, 3,250, …, 30,000}.

### Image generation

Our image generation process closely follows the approach described in Arnab et al. [2025]. Unlike the original framework in Arnab et al. [2025], we did not perform image resizing, as our approach did not require adapting images to fit predefined neural network input sizes. We omitted this step while retaining the core methodology.

For each simulated replicate, we retained only bi-allelic single nucleotide polymorphisms (SNPs) with a minor allele count of at least three (*i.e.*, singletons and doubletons were removed). We then generated a matrix **M** of dimension *n* × 499, where *n* represents the number of haplotypes in a replicate. Each row of **M** corresponds to a sampled haplotype, whereas each column represents one of the 499 SNPs. These SNPs were selected as follows: the SNP closest to the central position, the 249 closest SNPs upstream of this central SNP, and the 249 closest SNPs downstream of this central SNP. The central position in a simulated genomic region is the site that beneficial mutations are introduced in sweep simulations. The element **M***_ij_* takes a value of zero if the haplotype in the *i*th row contains the major allele at the SNP in the *j*th column and a value of one if it contains the minor allele.

To create a structured representation of genomic variation for improved pattern recognition, we constructed a transformed matrix **X** of dimension *n* × 237. This transformation was achieved by applying a sliding window approach over the columns of **M**, using windows of 25 SNPs with a stride of two between each window. Specifically, for *j* ∈ {1, 2, …, 237}, we computed the minor allele counts for each haplotype within a genomic window of length 25 SNPs, starting at column 2*j* − 1 of **M**. These minor allele count values were then sorted in increasing order and assigned to column *j* of **X**. As a result, rows toward the top of **X** represent haplotypes with a greater number of major alleles, whereas rows toward the bottom contain haplotypes with more minor alleles. This transformation provides a condensed summary of minor allele distributions across haplotypes in a structured manner, enhancing downstream image-based learning approaches.

### Histogram of oriented gradients

Histogram of oriented gradients (HOG) is a feature descriptor for object detection and image analysis, capturing local gradient information by computing histograms of gradient orientations within small spatial regions of an image [Nixon and Aguado, 2019, Szeliski, 2022]. Given an im-age matrix **X** ∈ R*^n^*^×237^ (see *Image generation* subsection above), the gradient is computed using finite differences. At pixel (*i, j*), the horizontal gradient is defined as *G_x_*(*i, j*) = **X***_i_*_+1_*_,j_* − **X***_i_*_−1_*_,j_* for *i* ∈ {2, 3, 4 *…, n* − 1} and *j* ∈ {1, 2, 3 *…,* 237}, and the vertical gradient is defined as *G_y_*(*i, j*) = **X***_i,j_*_+1_ − **X***_i,j_*_−1_ for *i* ∈ {1, 2, 3 *…, n*} and *j* ∈ {2, 3, 4 *…,* 236}. From these gradients, the magnitude and orientation at pixel (*i, j*) are respectively obtained as [Dalal and Triggs, 2005]

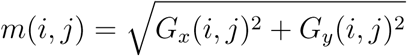

and

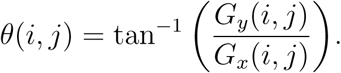

Each pixel (*i, j*) thus provides a vector [*m*(*i, j*), *θ*(*i, j*)] indicating a local gradient strength and direction. The image is then divided into small spatial regions termed cells [Dalal and Triggs, 2005], where an orientation histogram is formed by voting the pixel gradients into a fixed number of orientation bins. Each pixel casts a vote to the bin corresponding to its *θ*, with a weight equal to its gradient magnitude *m*.

To enhance robustness to illumination changes, HOG normalizes these histograms using larger overlapping regions by grouping several cells into blocks [Dalal and Triggs, 2005]. Each block gathers the cell histograms within it into a single vector *v*. This vector is then normalized by applying the “L2-Hys” scheme [Dalal and Triggs, 2005], which first applies *L*_2_ normalization of *v* to yield 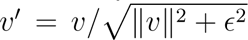, where *ɛ* is a small constant to prevent division by zero, followed by component-wise clipping that ensures each element of *v*^′^ exceeding 0.2 is capped at 0.2 [Dalal and Triggs, 2005]. This strategy was reported to achieve superior detection accuracy compared to regular *L*_2_ normalization in the original formulation of HOG [Dalal and Triggs, 2005]. The normalized descriptor for each block is treated as one HOG descriptor vector, and the final HOG feature is the concatenation of all block vectors from the dense, overlapping grid covering the image. We implemented this technique using the skimage.feature.hog function from the scikit-image Python library.

### Positive-unlabeled learning algorithm

In PU learning, we can only make use of labeled positive samples and unlabeled samples, where the unlabeled set may contain both positive and negative samples. In our context, if positive samples represent sweep regions, then negative samples would be non-sweep regions, with the collection of regions of the genome (unlabeled set) a mixture of sweep and non-sweep regions. The goal of the PU learning algorithm is to estimate the probability of a sample belonging to the positive class despite the absence of explicitly labeled negative samples [Elkan and Noto, 2008]. Let *X* ∈ R*^p^* denote an input vector for a genomic region and *Y* ∈ {0, 1} denote the true class label, where *Y* = 1 represents a positive sample and *Y* = 0 represents a negative sample. In this framework, a subset of the positive samples are chosen to be labeled, which implies that every labeled sample is positive, but not every positive sample is labeled. We introduce the label state *S* ∈ {0, 1} to denote whether a sample is (*S* = 1) or is not (*S* = 0) labeled. We assume the standard selected completely at random (SCAR) condition [Kumar and Lambert, 2024] that the chance of a positive being labeled is independent of its feature values. That is, positives are randomly selected for labeling from the set of all positives, implying that the labeling probability for any given positive sample is a constant *c*. Under the SCAR assumption, *P* (*S* = 1|*Y* = 1*, X*) = *P* (*S* = 1|*Y* = 1) = *c*. This constant *c*, termed the label frequency or propensity [Kumar and Lambert, 2024], represents the probability that a true positive is included among the labeled training samples.

Though traditional classifiers compute the conditional probability *P* (*Y* |*X*) of the class *Y* given input vector *X*, for PU learning, we construct a “nontraditional classifier” that computes the conditional probability *P* (*S*|*X*) of the sample label state *S* given the input vector *X* [Elkan and Noto, 2008]. In particular, whereas the traditional binary classifier estimates the quantity *f* (*X*) = *P* (*Y* = 1|*X*), we train a model on the available labeled positives (as positive class) and unlabeled data (treated as negatives) to estimate

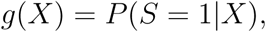

which represents the probability that a sample is labeled [Elkan and Noto, 2008]. This function *g*(*X*) is not the same as the true class probability *f* (*X*) = *P* (*Y* = 1|*X*), but the two are closely related. Given that *S* = 1 can only occur for positive samples, *i.e.*, *P* (*S* = 1|*Y* = 0*, X*) = 0, and that the label frequency under SCAR is *P* (*S* = 1|*Y* = 1*, X*) = *c* [Elkan and Noto, 2008], we have by the law of total probability that

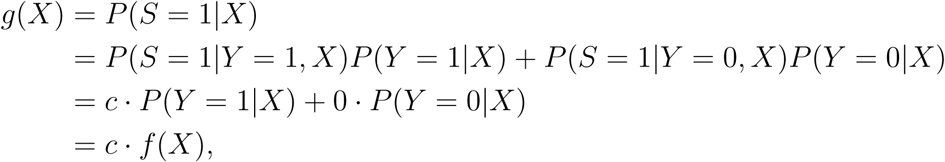

and thus *f* (*X*) = *g*(*X*)*/c*. This result allows us to transform the nontraditional classifier into a traditional classifier, provided that we can estimate *c*. To estimate *c* from the labeled and unlabeled samples, we hold out a small subset of the known positive samples as a validation set V. After training the classifier *g*(*X*) on the remaining positive and unlabeled samples, we apply it to the held-out positive samples in V to estimate the labeling probability. An unbiased estimator for *c* is the mean predicted probability on this held-out set, computed as

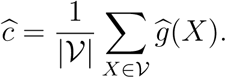

Because *g*(*X*) estimates *P* (*S* = 1|*X*), its mean over a representative set of actual positives approximates the fraction of positives that are labeled. Using this *c* value, we then correct the output of the model outputs to compute the final PU classifier

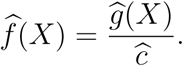

Here *f* (*X*) is our estimate of *P* (*Y* = 1|*X*) for each sample, which for our study is the probability that a given genomic region is truly a sweep. We implemented this two-step PU learning approach using the *ElkanotoPuClassifier* module from the *pulearn* library, which wraps the Elkan and Noto algorithm [Elkan and Noto, 2008] in a *scikit-learn* compatible estimator like *LogisticRegression*. All predicted probabilities were corrected by the factor *c*, and the resultant *f* (*X*) values were used for downstream classification of sweep versus non-sweep regions.

### Choosing the best feature extraction pipeline

To determine the optimal feature extraction pipeline for a downstream PU learning architecture, we evaluated different preprocessing techniques, HOG parameters, and post-processing methods. This process of choosing the best pipeline ensures that extracted features effectively capture distinguishing patterns. We evaluated the feature extraction pipelines under both *CEU-CEU* and *Constant-CEU* scenarios. To keep pipeline selection simple, we chose the PU learning model base classifier as logistic regression with no penalty, ensuring that model selection focuses purely on feature generation. We also opted for 10,000 labeled sweeps and 10,000 unlabeled samples for both scenarios. In both cases, we maintained a consistent imbalanced ratio of sweep and neutral replicates in the unlabeled set, with 1,000 sweep replicates (10% of the unlabeled set). The effect of additional imbalance ratios, as well as hyperparameter tuning for the base classifier are assessed in the *Model performance as function of unlabeled set class imbalance* and *Improving detection performance with model calibration* subsections of the *Results*, respectively.

We considered preprocessing input images before computing HOG features to aid in providing consistency across images. We tested four approaches: no preprocessing, histogram equalization [Gonzalez, 2009], image normalization (scaling pixel values in an image to range from zero to one), and division by standard deviation (scaling pixel values in an image so that its pixel values have a standard deviation of one). While no preprocessing serves as a baseline, histogram equalization improves contrast by redistributing pixel intensities, whereas both image normalization and division by standard deviation reduces brightness differences among images while preserving relative intensity information within images.

After preprocessing, we extracted features using HOG with different parameter settings, chosen based on prior studies [Lowe, 2004a, Dalal and Triggs, 2005]. The working principle of HOG is detailed in the *Histogram of oriented gradients* subsection above. We varied the number of gradient orientations (six, nine, or 12), pixel sizes per cell (8 × 8 or 16 × 16), and cells per block (2 × 2 or 3 × 3) to determine which configuration best captures local patterns for better discrimination between sweep and non-sweep. These choices impact the level of detail retained in the feature representation, with smaller cells capturing finer details and larger cells providing coarser patterns.

We applied post-processing to the extracted HOG features to further refine the input for the PU learning model. We tested no post-processing, sample-wise mean subtraction (ensuring a mean of zero across the features of a single sample), feature-wise standardization (ensuring a mean of zero and standard deviation of one across samples), and sample-wise normalization (ensuring feature values from zero to one across a single sample). These transformations help mitigate biases in feature distributions and improve comparability across different images.

Based on AUPRC scores, we found that two post-processing approaches, feature-wise standardization and sample-wise normalization, were the least effective in both *CEU-CEU* and *Constant-CEU* scenarios (Tables S7 and S8). Consequently, we excluded these methods from further analysis. In the *CEU-CEU* scenario, the highest AUPRC of 0.7327 was achieved using a feature extraction pipeline with no preprocessing, six gradient orientations, a pixel size of 16 × 16 per cell, a block size of 3 × 3 cells, and sample-wise mean subtraction as post-processing. The second-highest AUPRC of 0.7326 was obtained with the same configuration, but without any post-processing. Given that these two pipelines performed nearly identically, we selected the second configuration due to its comparative simplicity. We refer to this pipeline as *P1* going forward.

In the *Constant-CEU* scenario, the overall performance was noticeably poorer compared to the *CEU-CEU* scenario, as expected. The best AUPRC achieved was 0.3544 using a pipeline with histogram equalization as preprocessing, nine gradient orientations, a pixel size of 16 × 16 per cell, a block size of 3 × 3 cells, and no post-processing. The second-best AUPRC was 0.3522, obtained with the same configuration but incorporating sample-wise mean subtraction as post-processing. Given the marginally higher performance and the reduced complexity of the best-performing configuration, we selected the first one as the optimal pipeline for this scenario, which we denote as the *P2* pipeline.

Comparing these results to the *CEU-CEU* scenario, we observe that the inclusion of histogram equalization and a higher number of gradient orientations (nine instead of six) led to improved performance in the *Constant-CEU* setting. Histogram equalization enhances contrast by redistributing pixel intensities, which likely helps mitigate variation in image brightness and improves edge detection in a dataset with more heterogeneous samples. Similarly, increasing the number of gradient orientations allows the HOG descriptor to capture more detailed directional patterns, which may be particularly beneficial when dealing with the greater domain mismatch in the *Constant-CEU* scenario.

To assess the generalizability of the selected pipelines, we compared the performance of *PULSe* across both scenarios using the *P1* and *P2* pipelines, and respectively denote these method variants by *PULSe*[*P1*] and *PULSe*[*P2*]. When evaluated on the *CEU-CEU* scenario, *PULSe*[*P2*] exhibited a slight decline in performance, with an AUPRC drop of 0.014 and an MCC drop of 0.0124 compared to *PULSe*[*P1*] (Figure S6). Conversely, in the *Constant-CEU* scenario, *PULSe*[*P1*] demonstrated a larger AUPRC decline of 0.0274 relative to *PULSe*[*P2*], and an MCC decline of 0.0142 (Figure S6). These results suggest that the *P2* pipeline offers slightly stronger performance in case of domain mismatch, while the *P1* pipeline showcases improved performance in case of domain match scenario.

### Training the *smbCNN_GRL*[*PI*] architecture

To train the *smbCNN GRL*[*PI*] model under the *Constant–CEU* scenario, we assembled a training set consisting of 9,000 sweep and 9,000 neutral replicates each from both CEU and constant population size demographic histories. The model jointly learned to classify sweeps and neutral samples while also determining the source domain of each input, requiring two forms of supervision: class labels and domain labels. The constant population size training replicates were annotated with both class labels (sweep or neutral) and a domain label of zero. In contrast, the CEU replicates were only assigned a domain label of one, and their class labels were masked during training, meaning they were excluded from the classification loss calculation. Construction of the validation set followed a similar procedure, using 1,000 sweep and 1,000 neutral replicates each from the CEU and constant population size histories. Early stopping [Prechelt, 2002] based on lowest validation loss was used to prevent overfitting. An identical training strategy was applied for the *CEU–CEU* scenario, except that all replicates (both domain label zero and one) were drawn from the CEU population. No constant population size replicates were used in this test, allowing us to evaluate domain-invariant performance in a demographically matched scenario. The test set used for evaluation was identical to the unlabeled samples employed in both *PULSe* models, enabling a consistent comparison across models.

### Empirical data filtration and image generation

To apply *PULSe* on empirical data, we used phased haplotype data from the 99 CEU and 86 BEB individuals of the 1000 Genomes Project dataset [1000 Genomes Project Consortium, 2015]. Consistent with our simulation protocol, we retained only biallelic SNPs with a minor allele count of at least three, thereby removing singletons and doubletons. To mitigate the potential influence of technical artifacts, we followed the filtering protocol of Mughal et al. [2020] and excluded genomic segments of length 100 kilobases with mean CRG (Consensus Reference Genomes) mappability and alignability scores below 0.9 [Talkowski et al., 2011]. These scores reflect the confidence with which sequencing reads can be accurately mapped to specific genomic regions, and their use ensures that downstream analyses are restricted to reliably interpretable regions of the genome.

We then applied the *PULSe* image generation approach described in the *Image Generation* subsection to all 22 autosomes. For each autosome, we began with the first 499 SNPs and generated the image representation of the spatial change in genomic variation using the same transformation pipeline used for simulated samples. Specifically, the haplotype matrix across these 499 SNPs was converted into a structured *n* × 237 matrix that summarizes haplotypic variation across overlapping genomic windows, where *n* = 198 is the number of haplotypes from the CEU sample and *n* = 172 is the number of haplotypes from the BEB sample. We then advanced the SNP window by 50 SNPs and repeated the process until reaching the end of the chromosome. Each image was assigned a genomic coordinate corresponding to the 250th SNP within the 499-SNP window. This approach yielded a dense, genomewide series of *PULSe*-compatible images that retain biologically meaningful haplotype structure and are suitable for inference using our trained models.

## Acknowledgments

This work was supported by National Institutes of Health grant R35GM128590, by National Science Foundation grants DBI-2130666 and DEB-1949268, and by the UKRI Natural Environment Research Council grant NE/Y003519/1. Computations for this research were performed using the services provided by Research Computing at Florida Atlantic University.

## Data Availability

We release the source code for *PULSe* under the MIT open source license, and this repository can be accessed on GitHub (https://github.com/sandipanpaul06/PULSe/). The CEU and BEB data from the 1,000 Genomes Project can be accessed from the project website (https://www.internationalgenome.org/category/phase-3/).

