## Supplemental Figures and Tables for "Semi-supervised detection of natural selection with positive-unlabeled learning"

### Supplementary material

Table S1: Gene Ontology (GO) Biological Process for CEU candidate genes identified by *PULSe* compared against a background set of all retained genes based on *PULSe* calibrated prediction output. See file TableS1.csv.

Table S2: Gene Ontology (GO) Cellular Component for CEU candidate genes identified by *PULSe* compared against a background set of all retained genes based on *PULSe* calibrated prediction output. See file TableS2.csv.

Table S3: Gene Ontology (GO) Molecular Function for CEU candidate genes identified by *PULSe* compared against a background set of all retained genes based on *PULSe* calibrated prediction output. See file TableS3.csv.

Table S4: Gene Ontology (GO) Biological Process for BEB candidate genes identified by *PULSe* compared against a background set of all retained genes based on *PULSe* calibrated prediction output. See file TableS4.csv.

Table S5: Gene Ontology (GO) Cellular Component for BEB candidate genes identified by *PULSe* compared against a background set of all retained genes based on *PULSe* calibrated prediction output. See file TableS5.csv.

Table S6: Gene Ontology (GO) Molecular Function for BEB candidate genes identified by *PULSe* compared against a background set of all retained genes based on *PULSe* calibrated prediction output. See file TableS6.csv.

Table S7: Gain in *PULSe*[*P1*] area under the precision-recall curve (AUPRC) due to the application of each pre-processing, post-processing, and histogram of oriented gradients (HOG) parameter choice, measured by holding all other modeling components at their default: no-preprocessing, nine orientations, $8 \times 8$ pixels per cell, $3 \times 3$ cells per block, and no-postprocessing.

| Category | Technique | AUPRC gain |
|---|---|---|
| Pre-processing | Histogram equalization | +0.003 |
| | Image normalization | −0.021 |
| | Division by standard deviation | −0.048 |
| HOG Parameters | Number of gradient orientations | +0.112 |
| | Pixel sizes per cell | +0.108 |
| | Cells per block | +0.006 |
| Post-processing | Sample-wise mean subtraction | +0.029 |
| | Feature-wise standardization | −0.001 |
| | Sample-wise normalization | +0.001 |

Table S8: Gain in $PULSe[P2]$ area under the precision-recall curve (AUPRC) due to the application of each pre-processing, post-processing, and histogram of oriented gradients (HOG) parameter choice, measured by holding all other modeling components at their default: no-preprocessing, nine orientations, $8 \times 8$ pixels per cell, $3 \times 3$ cells per block, and no-postprocessing.

| Category | Technique | AUPRC gain |
|---|---|---|
| Pre-processing | Histogram equalization | +0.066 |
| | Image normalization | −0.001 |
| | Division by standard deviation | −0.003 |
| HOG Parameters | Number of gradient orientations | +0.071 |
| | Pixel sizes per cell | +0.053 |
| | Cells per block | +0.036 |
| Post-processing | Sample-wise mean subtraction | +0.007 |
| | Feature-wise standardization | +0.001 |
| | Sample-wise normalization | +0.001 |

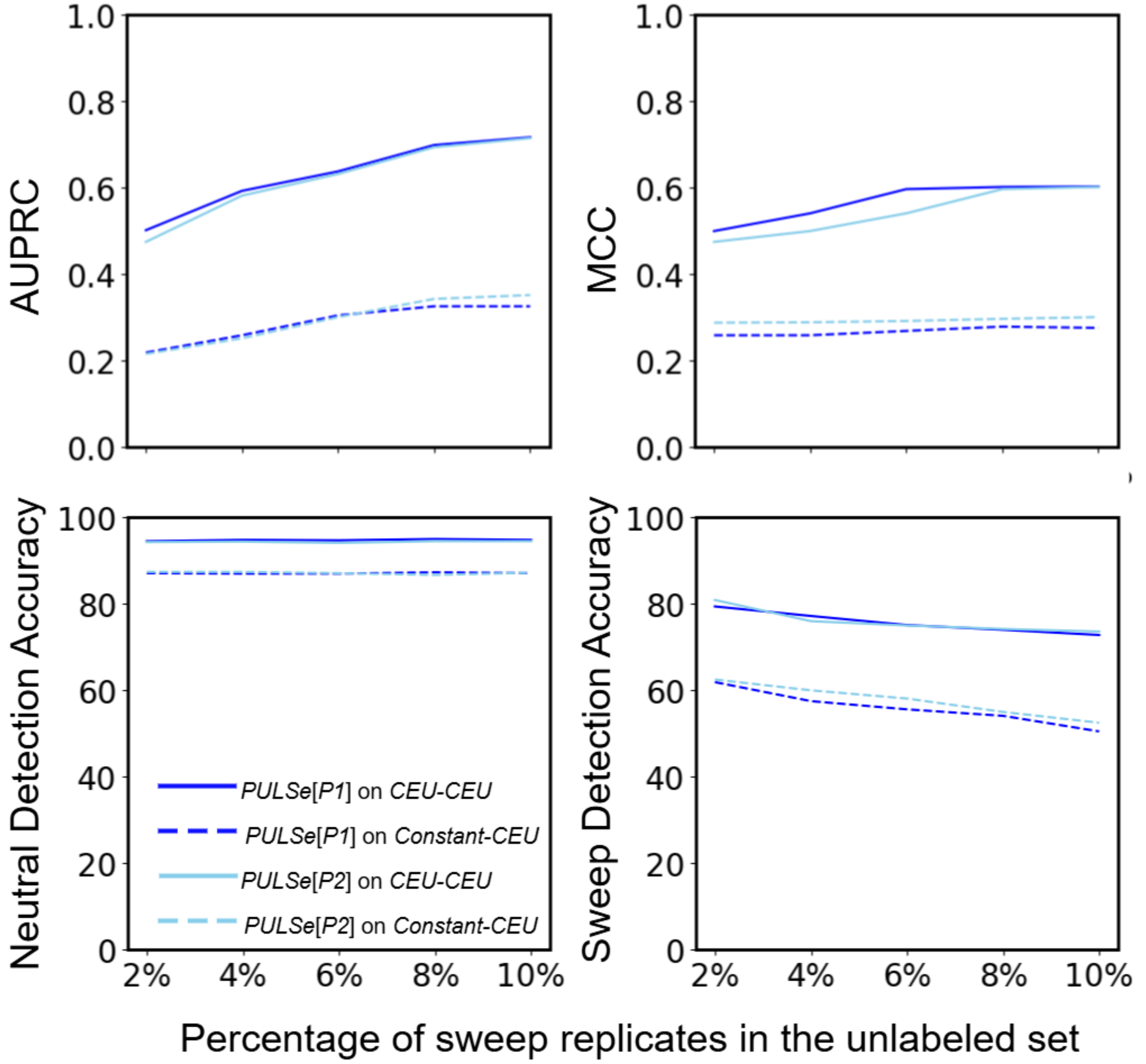


Figure S1: Change in area under the precision-recall curve (AUPRC), the Matthews correlation coefficient (MCC), neutral detection accuracy, and sweep detection accuracy of *PULSe*[*P1*] and *PULSe*[*P2*] for both *CEU-CEU* and *Constant-CEU* scenarios as a function of the percentage of sweep replicates in the unlabeled sets. The percentage of sweep replicates ranges from 2% to 10% in increments of 2%, with the total number of both labeled and unlabeled samples fixed at 10,000.

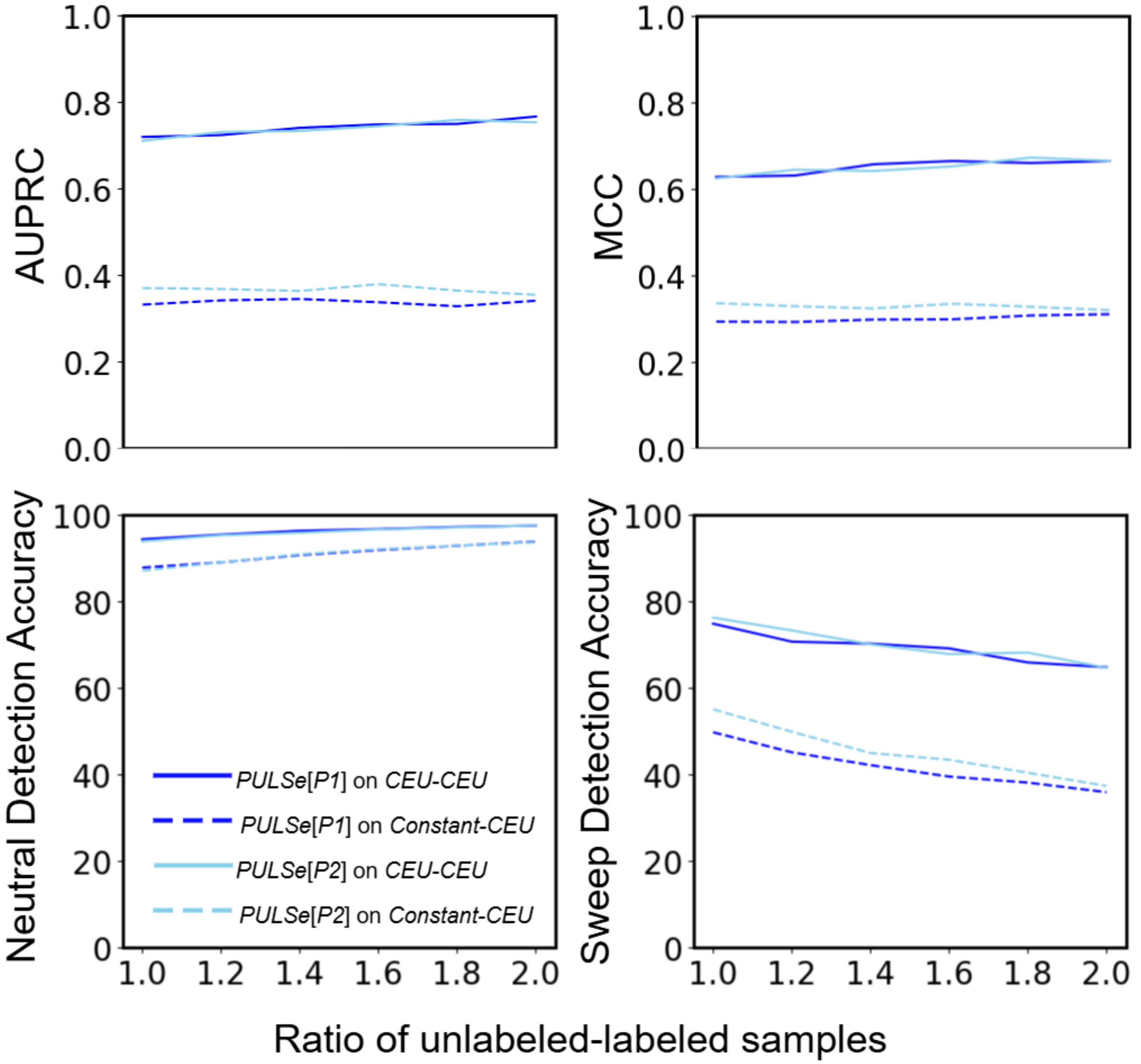


Figure S2: Change in area under the precision-recall curve (AUPRC), the Matthews correlation coefficient (MCC), neutral detection accuracy, and sweep detection accuracy of *PULSe*[*P1*] and *PULSe*[*P2*] for both *CEU-CEU* and *Constant-CEU* scenarios as a function of the ratio of unlabeled to labeled set sizes. The ratio of unlabeled to labeled set sizes ranges from 1.0 to 2.0 in increments of 0.2, with the total number of labeled samples fixed at 10,000.

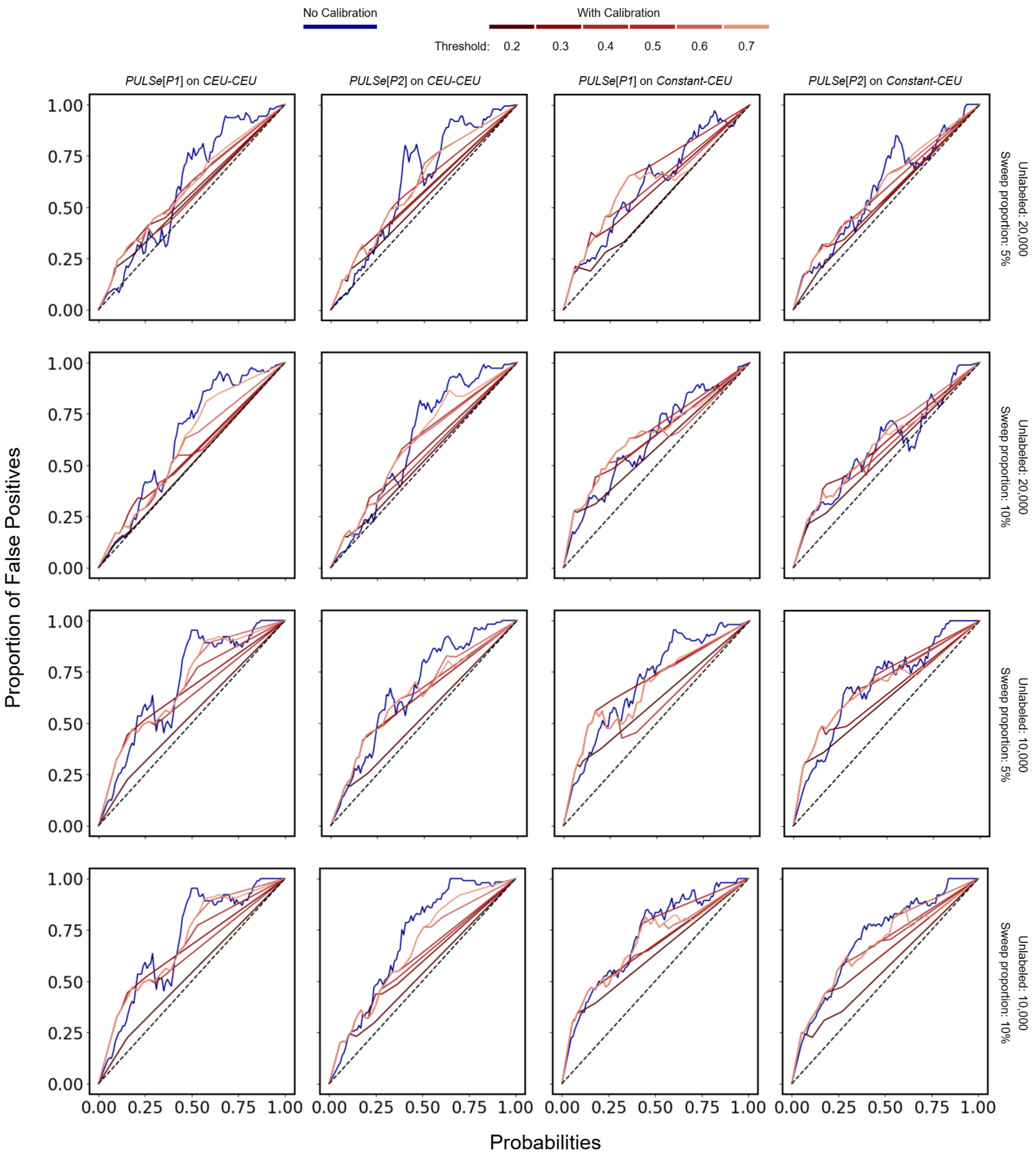


Figure S3: Calibration curves of calibrated (see *Improving detection performance with model calibration* subsection of the *Results* for details) *PULSe*[*P1*] and *PULSe*[*P2*] models for both *CEU-CEU* and *Constant-CEU* scenarios using calibration thresholds ranging from 0.2 to 0.7 in increments of 0.1, compared against calibration curves of uncalibrated *PULSe*[*P1*] and *PULSe*[*P2*] models. The total number of labeled samples was fixed at 10,000. The composition of the unlabeled set was varied in two ways: the number of unlabeled samples (10,000 or 20,000) and the percentage of sweep samples in the unlabeled set (5% or 10%).

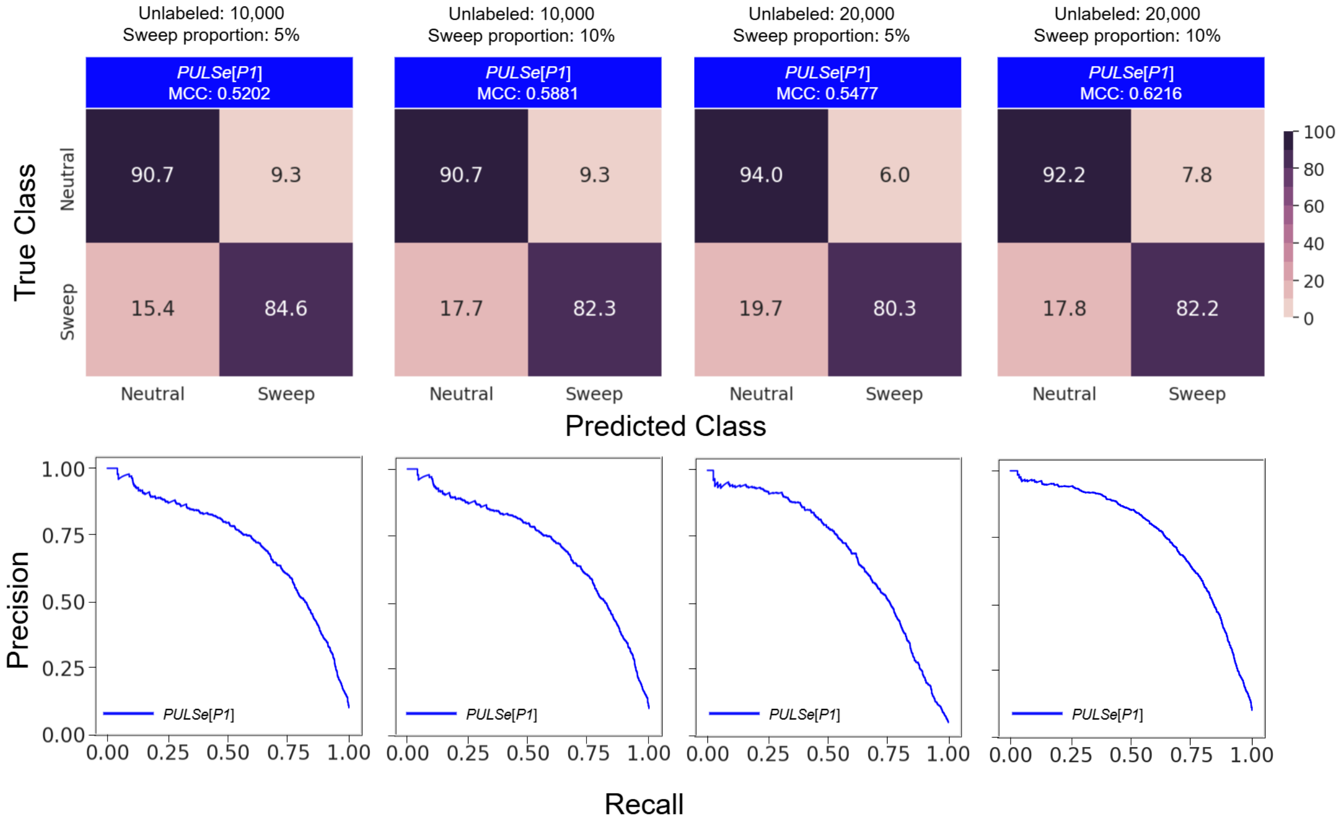


Figure S4: Classification rates and accuracies as depicted by confusion matrices (top row) and reliability to detect sweeps as depicted by precision-recall curves to differentiate sweeps from neutrality (bottom row) on the unlabeled set for the *CEU-CEU* scenario (see *Modeling Description* subsection of the *Results*) for the calibrated *PULSe*[*P1*] model using a calibration threshold of 0.3. The composition of the unlabeled set was varied in two ways: the number of unlabeled samples (10,000 or 20,000) and the percentage of sweep samples in the unlabeled set (5% or 10%).

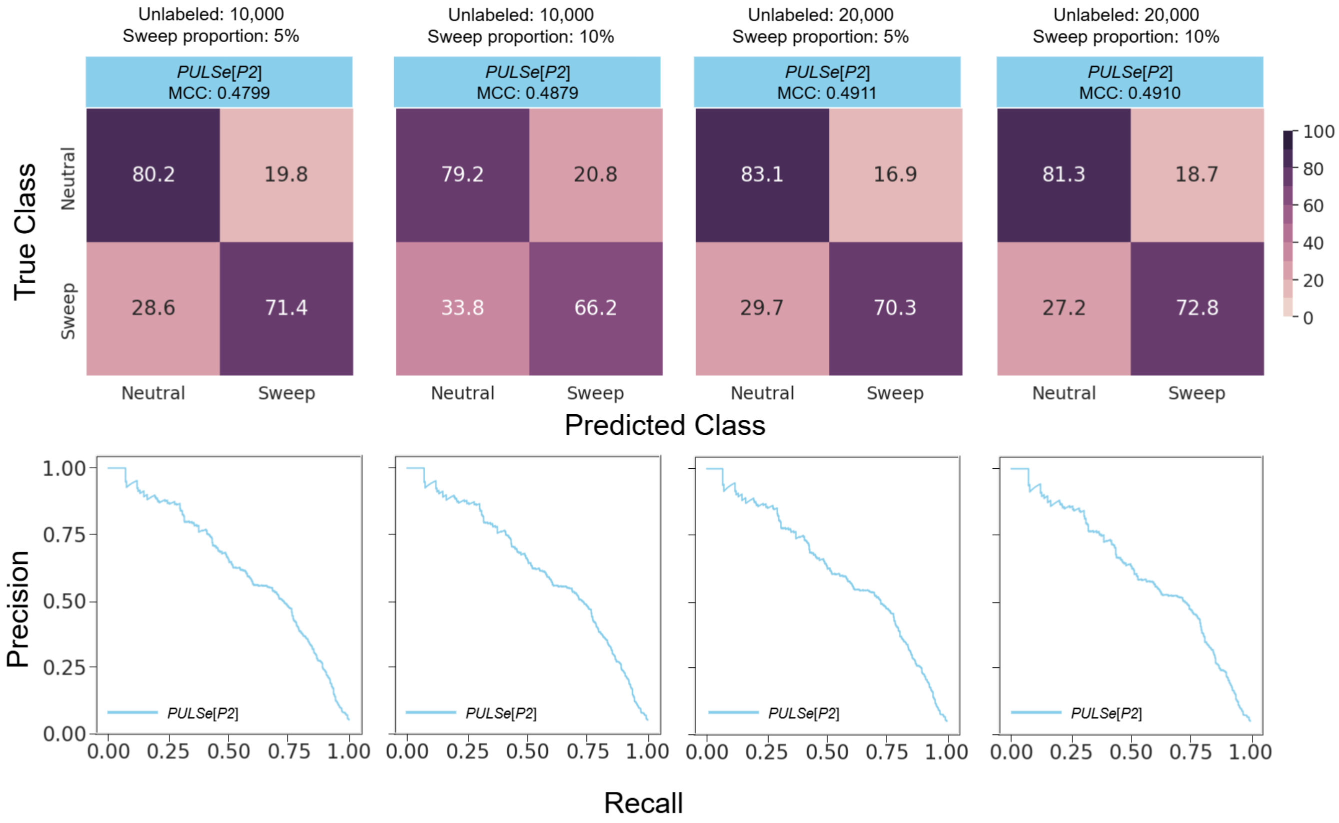


Figure S5: Classification rates and accuracies as depicted by confusion matrices (top row) and reliability to detect sweeps as depicted by precision-recall curves to differentiate sweeps from neutrality (bottom row) on the unlabeled set for the *Constant-CEU* scenario (see *Modeling Description* subsection of the *Results*) for the calibrated *PULSe*[*P2*] model using a calibration threshold of 0.3. The composition of the unlabeled set was varied in two ways: the number of unlabeled samples (10,000 or 20,000) and the percentage of sweep samples in the unlabeled set (5% or 10%).

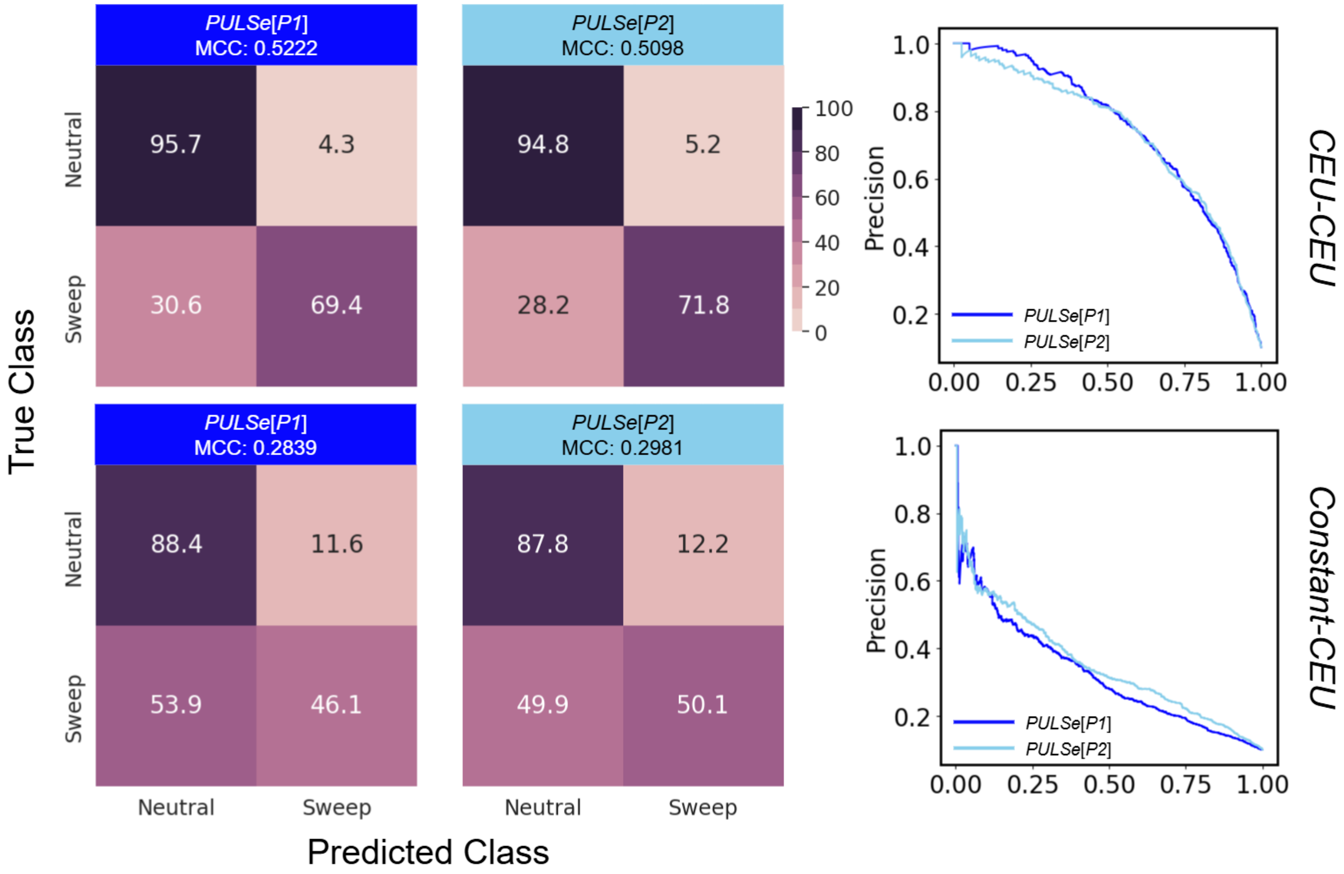


Figure S6: Classification rates and accuracies as depicted by confusion matrices and reliability to detect sweeps as depicted by precision-recall curves to differentiate sweeps from neutrality on the unlabeled set for the *CEU-CEU* (top row) and *Constant-CEU* (bottom row) scenarios (see *Modeling Description* subsection of the *Results*) for the *PULSe*[*P1*] and *PULSe*[*P2*] models. For both scenarios, the unlabeled set contains 9,000 neutral and 1,000 sweep replicates simulated based on the recent strong bottleneck demographic history of the CEU human population from the 1000 Genomes Project dataset.

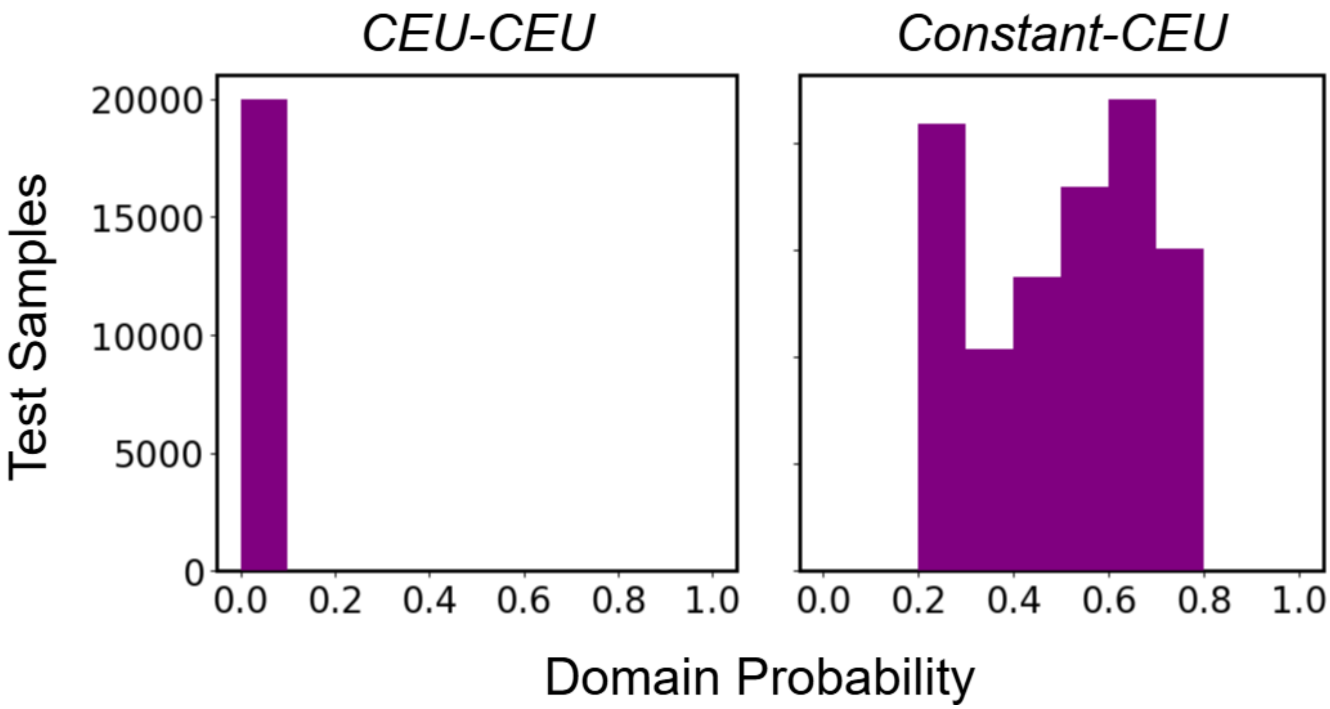


Figure S7: Histograms summarizing the predicted domain probability distributions of *smbCNN_GRL[PI]* for the *CEU-CEU* (left) and *Constant-CEU* (right) scenarios. Predicted domain probabilities reflect how confidently the domain classifier within *smbCNN_GRL[PI]* assigns unlabeled samples to in-domain (probability $< 0.5$) versus out-of-domain (probability $> 0.5$) distributions.

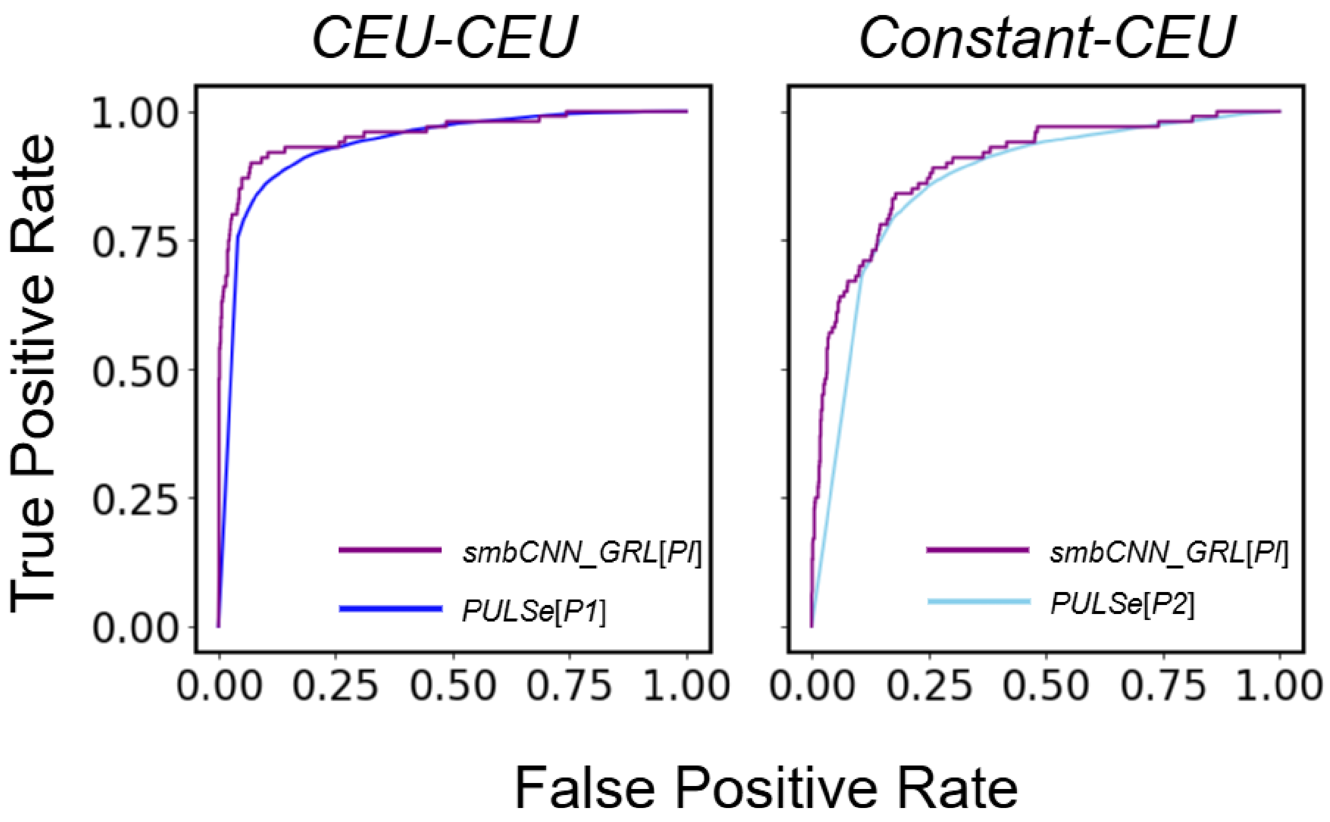


Figure S8: Powers (true positive rates) to detect sweeps as depicted by receiver operating characteristic curves to differentiate sweeps from neutrality on the *CEU-CEU* (left) and *Constant-CEU* (right) scenarios for *smbCNN_GRL*[*PI*] and *PULSe* models.
